# Sweetwater: an underrated crude glycerol for sustainable lipid production in non-conventional yeasts

**DOI:** 10.1101/2023.09.18.558176

**Authors:** Valériane Malika Keita, Yi Qing Lee, Meiyappan Lakshmanan, Dave Siak-Wei Ow, Paul Staniland, Jessica Staniland, Ian Savill, Kang Lan Tee, Tuck Seng Wong, Dong-Yup Lee

## Abstract

Sweetwater, a methanol-free by-product of the fat splitting process, is an emerging alternative feedstock for microbial utilization of crude glycerol. Yeasts are promising candidates in this context due to their versatility in delivering a wide range of value-added products including lipids. To enhance sweetwater utilization, we investigated the growth and lipid production of 21 oleaginous yeast strains in sweetwater and found that nutrient limitation and the unique carbon composition of sweetwater boosted significant lipid accumulation in several strains, in particular *Rhodosporidium toruloides* NRRL Y-6987. To decipher the underlying mechanism, the transcriptomic changes of *R. toluroides* NRRL Y-6987 were further analyzed, indicating potential sugars and oligopeptides in sweetwater supporting growth and lipid accumulation as well as exogenous fatty acid uptake leading to the enhanced lipid accumulation. Our comparative study successfully demonstrated sweetwater as a cost-effective feedstock and suggested potential sweetwater type and strain engineering targets for increasing microbial lipid production.

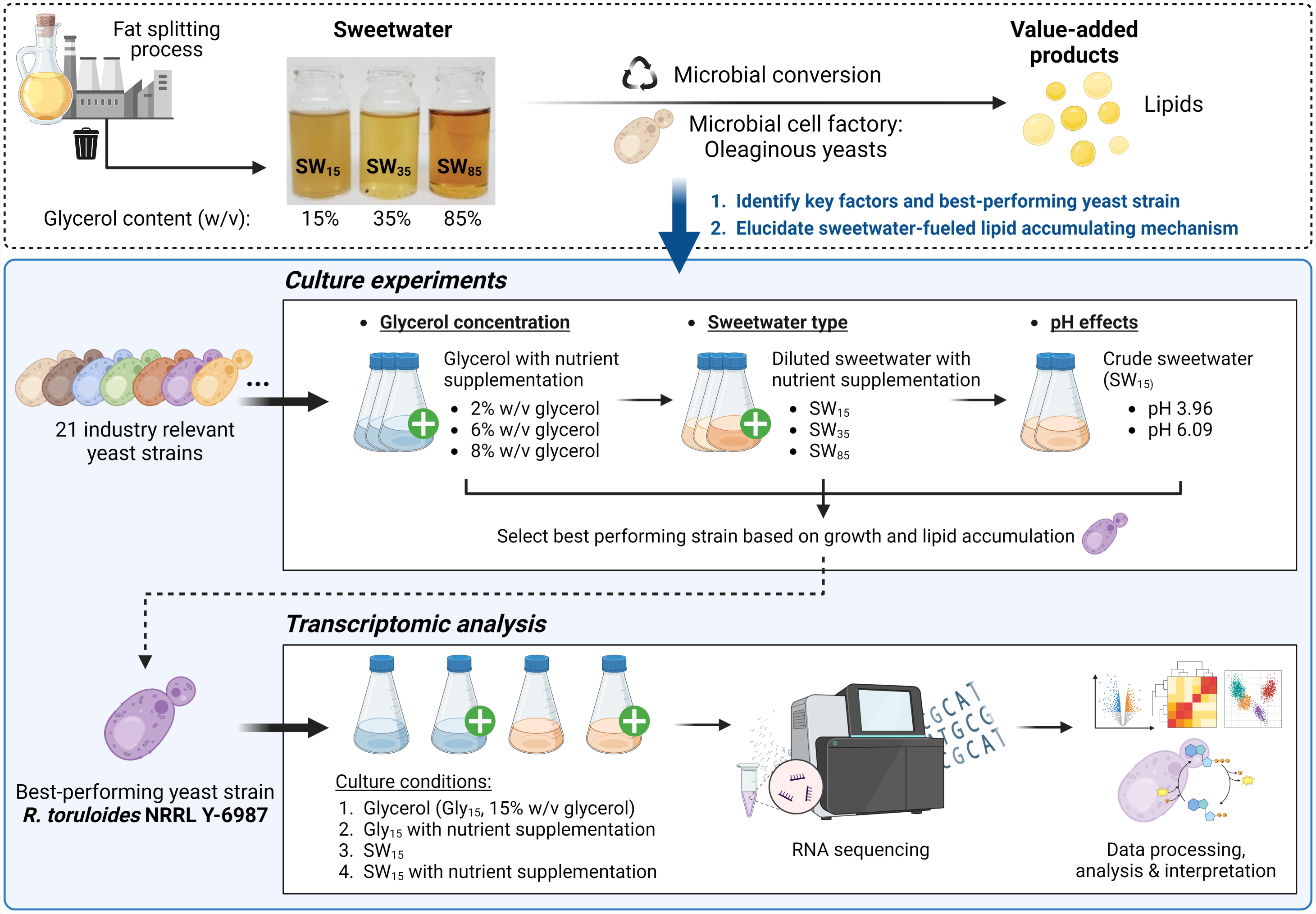

## 1. Introduction

The rapid expansion of the biodiesel industry has flooded the market with high volumes of its by-product, crude glycerol. This endangers the economic viability of biodiesel production and raises a disposal issue. To tackle this challenge, microbial conversion of crude glycerol into value-added chemicals has received a lot of interest due to its alignment with current expectations for sustainability and renewability. In fact, both chemical and biological conversions of crude glycerol are currently under investigation (Kaur *et al*., 2020; Luo *et al*., 2016). Notably, the ability of several microorganisms to grow on crude glycerol has been evaluated, and led to the successful production of various chemicals such as polyols, polyhydroxyalkanoates, organic acids, biogas, biofuels, proteins of industrial relevance, glycerol derivatives and lipids (González-Villanueva *et al*., 2019). Among these potential products, lipids are particularly interesting because they could fit into a circular economy (Tomás-Pejó *et al*., 2021). For microbial lipid production, yeasts are notable for their genetic adaptability, versatile carbon utilization, short duplication times and high lipid accumulation potential (Thevenieau and Nicaud, 2013). Despite extensive research on yeast lipid production, current yields remain suboptimal due to inhibitory effects of methanol (Gao *et al*., 2016; Samul *et al*., 2014). Furthermore, elevated glycerol concentrations also negatively impact cell growth (Muniraj *et al*., 2015). To overcome these limitations, crude glycerol can be diluted to lower the concentration of impurities and glycerol (Ito *et al*., 2005). Alternatively, feeding strategies during microbial fermentation can be manipulated to increase conversion efficiency (Signori *et al*., 2016).

There has been a growing interest in harnessing alternative sources of crude glycerol discharged from fat splitting and saponification processes, which contribute to approximately 30% and 6% of the global glycerol production, respectively (Ciriminna *et al*., 2014; Pagliaro, 2017). Crude glycerol from the fat splitting process is widely referred to as sweetwater. Its methanol-free composition suggest that it might be a more compatible feedstock for lipid production than biodiesel-derived crude glycerol. Furthermore, sweetwater is available in different purity levels: SW_15_ which has glycerol concentration of 15% (w/v) with a low pH and high brownish fatty residue content (**Fig. S1**), as well as cleaner forms with glycerol concentrations of 35% (w/v) (SW_35_) and 85% (w/v) (SW_85_). SW_85_ is considered a semi-crude glycerol and has low economic value of around 0.04 – 0.09 USD/per pound (Mota *et al*., 2017). Therefore, sweetwater could be more advantageous than biodiesel-derived crude glycerol as a microbial lipid production feedstock in terms of efficiency and cost.

To enhance the utilization of sweetwater, we comprehensively evaluated the suitability of sweetwater as a feedstock using 21 selected oleaginous yeast species by comparing both cell growth and lipid accumulation across different glycerol and impurities concentrations, as well as media pH. We observed lipid production induced by sweetwater and its underlying mechanism was further elucidated through an in-depth analysis of the associated transcriptomic changes.

## 2. Materials and Methods

### 2.1 Media and sweetwater formulation

Actively growing cultures and pre-cultures were prepared using YM agar plates (containing 20 g/L agar, 3 g/L yeast extract, 3 g/L malt extract, 5 g/L peptone, and 10 g/L glucose) and YPD broth (containing 10 g/L yeast extract, 20 g/L peptone, and 20 g/L glucose). Glycerol-based defined media consist of the following components: 1.9 g/L Yeast Nitrogen Base (Formedium, CYN0501), 0.79 g/L complete supplement mixture (Formedium, DSC0019), an appropriate amount of ammonium sulfate to achieve an C/N molar ratio of 60, and glycerol at concentrations of 15, 20, 80, or 160 g/L, along with a separate 150 g/L glycerol medium prepared without additional nutrients, using ultrapure water and microbiology-grade glycerol. All media were sterilized by autoclaving prior to use.

Croda (Hull, UK) supplied batches of sweetwater. The most unrefined form of sweetwater is SW_15_, with a glycerol content of 15% (w/v) and pH of 3.9 ± 0.1 @ 20°C, primarily attributed to free fatty acids present in sweetwater. Additionally, it contains a significant quantity of brownish fatty residues, captured by filters during sweetwater clarification process (**Fig. S1**). SW_35_ also exhibits similar properties. When collected further down the glycerol refining process, sweetwater becomes purer but also more exhibits higher alkalinity, featuring a glycerol content of 85% (w/v). Solid residues were removed by filtration prior to sweetwater use to prevent interference with optical density readings. Filtered sweetwater was further steam sterilized to avoid microbial contamination. To evaluate the pH effect, 1M sodium hydroxide was added to SW_15_ to increase the pH to 6 ± 0.1 (@ 20°C). To prepare SW_1.5_SC medium, sterile SW_15_, SW_35_ or SW_85_ was diluted with 2× SC nutrient mix (3.8 g/L Yeast Nitrogen Base, 1.58 g/L complete supplement mixture and 1.44 g/L ammonium sulfate) and ultrapure water. SW_35_SC was prepared by directly dissolving 1.9 g/L Yeast Nitrogen Base, 0.79 g/L complete supplement mixture and 0.72 g/L ammonium sulfate in SW_35_. The media was then autoclaved.

### 2.2 Yeast strains, cultivation conditions and growth monitoring

Yeast strains employed in this study can be found in **Table 1**. Cells were cultivated in a sterile environment. Strains were streaked onto YM agar plates and allowed to grow at 25°C until colonies reached approximately 2 mm. Plates were then stored at 4°C for a month. To prepare inoculations, a single colony was transferred into an appropriate volume of YPD, followed by incubation at 25°C (210 rpm) until early saturation. To compare growth, an appropriate amount of inoculum and fresh media were transferred into 96-well clear U-bottom polypropylene microtiter plates (Thermo Fisher Scientific) in 4 replicates, aiming for an initial OD_600_ of 0.3. Subsequently, Inoculated plates were sealed and placed in Titramax 1000 (Heidolph) at 25°C and 1050 rpm for cultivation. To monitor the cultures, a SpectraMax M2e reader (Molecular Devices) was employed to measure absorbance of the cultures at 600 nm (OD_600_) at specified time intervals. Blank correction was applied to recorded OD_600_. Then, kinetic parameters were subsequently determined individually for each replicate using Growthcurver (Sprouffske and Wagner, 2016) and were used for statistical analysis. To compare over 2 conditions, ANOVA was first performed, followed by Tukey’s post-hoc test or t-test. For lipid and transcriptomic analysis, cells were washed twice and suspended in 50-mL centrifuge tubes containing 5 mL of medium or 250-mL flasks containing 50 mL of medium. Cultivation was initiated with OD_600_ of 0.3 and cultures were incubated at 25°C (210 rpm). OD_600_ was monitored using BioPhotometer Plus spectrometer (Eppendorf).

**Table 1.** Strains investigated in this study.

| No. | Strains (3-letter code) | ID |
| --- | --- | --- |
| 1 | *Barnettozyma californica* (bca) | NRRL Y-1680 |
| 2 | *Clavispora xylofermentans* (cxy) | BCC 30719 |
| 3 | *Cutaneotrichosporon curvatus* (ccu) | NRRL Y-1511 |
| 4 | *Cyberlindnera saturnus* (csa) | NRRL YB-4312 |
| 5 | *Lipomyces lipofer* (lli) | DSM 70305 |
| 6 | *Lipomyces starkeyi* (lst) | NRRL Y-1388 |
| 7 | *Lodderomyces elongisporus* (lel) | NRRL YB-4239 |
| 8 | *Metschnikowia pulcherrima* (mpu) | NRRL Y-5941-53 |
| 9 | *Pichia kudriavzevii* (pku) | NRRL Y-7551 |
| 10 | *Rhodotorula glutinis* (rgl) | NRRL Y-2502 |
| 11 | *Rhodotorula mucilaginosa* (rmu) | NRRL Y-17283 |
| 12 | *Rhodotorula toruloides* (rto) | NRRL Y-6987 |
| 13 | *Rhodotorula toruloides* (rto) | NRRL Y-1091 |
| 14 | *Scheffersomyces stipitis* (sst) | NRRL Y-7124 |
| 15 | *Scheffersomyces stipitis* (sst) | NRRL Y-11545 |
| 16 | *Scheffersomyces stipitis* (sst) | BCC 15191 |
| 17 | *Schwanniomyces occidentalis* (soc) | NRRL Y-2477 |
| 18 | *Torulaspora delbrueckii* (tde) | NRRL Y-866 |
| 19 | *Wickerhamomyces anomalus* (wan) | NRRL Y-366 |
| 20 | *Yarrowia lipolytica* (yli) | BCC 64401 |
| 21 | *Yarrowia lipolytica* (yli) | Y203A |

Note: NRRL strains were obtained from the Agricultural Research Service (ARS) culture collection (Beltsville, USA), DSM strains from the Deutsche Sammlung von Mikroorganismen und Zellkulturen (DMSZ, Braunschweig, Germany), and BCC strains were provided by Dr. Pornkamol Unrean from BIOTEC, Thailand (Unrean and Champreda, 2017).

### 2.3 Nile red staining

250 μL to 750 μL of cell culture (depending on the cell density) were collected at target time points. Cells were harvested and washed twice with PBS pH 7± 0.1 @ 20°C (1.23 g/L monobasic potassium phosphate, 0.416 g/L dibasic sodium phosphate, 8 g/L sodium chloride and 0.201 g/L potassium chloride). Washed pellets were stored at -80°C for further use. Frozen pellets were thawed on ice and resuspended in PBS to reach an OD_600_ of 5. 17 μL of freshly prepared DMSO:PBS (1:1) solution, 166 μL of cells suspensions (or PBS as blank) and 17 μL of freshly prepared Nile Red in acetone (60 μg/mL) were added to a black 96-well clear flat-bottom polystyrene microtiter plate (Grenier Bio-One). Four technical replicates were created for each sample and blank. Fluorescence was measured over a time course of 30 minutes, using excitation wavelength of 489 nm and emission wavelengths of 535 nm and 625 nm. Only fluorescence readings at 625 nm are presented. The fluorescence intensities were blank corrected and normalized with the OD_600_ of the cell suspension used. For time-course monitoring of lipid accumulation, average and standard deviations from technical replicates were depicted using GraphPad Prism. Statistical comparisons were done using ANOVA, and Tukey’s post-hoc test was performed when applicable.

### 2.4 Neutral lipids extraction and thin layer chromatography (TLC)

1. *R. toruloides* NRRL Y-6987 was cultivated in 250-mL flasks containing SW_15_ and SW_1.5_SC for 4 and 10 days. Cells from a 50-mL culture were then collected through centrifugation (12 minutes, 3000 g, 4°C), and the resulting pellets were stored at -80°C. Lipid extraction and TLC were performed according to previously described procedure (Sobus and Holmlund, 1976). Pellets were thawed and vortexed for 1 min. Subsequently, 775 mg from each pellet was transferred into a sterile 50-mL polypropylene centrifuge tube (Fisher Scientific). 3 mL of HCl (1 M) was added to each tube for cell hydrolysis. Mixtures were subjected to vortexing for 1 min before incubation (2-h) at 78°C using a dry bath (Starlab). Mixtures were vortexed once for 1 min every 30 min at room temperature during incubation. After hydrolysis, cellular material was transferred into separating funnels and water was added to reach a final volume of 10 ml, followed by addition of an equal volume of chloroform/methanol mixture (1:1). Funnels were vigorously shaken, and the upper phase was removed after phase separation. The lower phase was subsequently washed with 10 mL of 0.1% (w/v) NaCl. Resulting upper phase was again removed after phase separation, and 10 mL of MilliQ water was used to wash the lower phase. After phase separation, target fractions were moved into 10-mL glass vials and dried at 65°C for 2-3 h using a dry bath (Starlab). Dried extracts were resuspended in 500 µL of hexane and transferred into 5-mL dark glass vials and stored at –20°C. Five µL samples were blotted onto TLC silica gel 60 F_254_ plates (Merck). Plates were developed in pre-saturated chambers using mobile phase of a mixture of hexane: diethyl ether: acetic acid (70:30:1). After development, plates were removed from the chamber and left to dry at room temperature. Spots were then developed by immersion in a *p*-anisaldehyde solution made from 300 mL of 95% (v/v) EtOH, 12 mL of *p*-anisaldehyde, 6 mL of glacial acetic acid and 12 mL of concentrated sulfuric acid, followed by heat gun drying. Erucic acid in EtOH (10 mg/mL) and a high erucic acid rapeseed (HEAR) oil splitting product in EtOH (20 mg/mL) were used as standards.

### 2.5 RNA extraction and sequencing

*R. toruloides* NRRL Y-6987 were grown in SW_15_, Gly_15_, SW_1.5_SC and Gly_1.5_SC. The first two media contained 15% (w/v) glycerol, and the latter two 1.5% (w/v) glycerol with nutrient supplementation. Three aliquots of 10 mL of culture at mid-exponential phase were harvested by centrifugation for 5 min. Cells were then suspended in 1 mL of RNA*later*^TM^ (Thermo Fisher Scientific) to stabilize the RNA, and mixed using a tube rotator for 1 h at 4°C. Cells were then centrifuged for 5 min, and resulting pellets were stored at -80°C. To dilute RNA*later* and aid cell sedimentation when necessary, 14 mL of sterile DEPC-treated ultrapure water was added. To ensure a RNAase-free environment, we utilized RNAase-free tips and tubes, and we decontaminated surfaces (including the biosafety cabinet, bottles, and pipettes) using RNaseZAP (Thermo Fisher Scientific) in accordance with the manufacturer’s guidelines. After thawing pellets on ice, they were resuspended in 1 mL of TRIzol reagent (Thermo Fisher Scientific). Suspensions were then moved into tubes with 250 μL of chilled, acid-washed glass beads (425-600 μm) (Sigma). Next, tubes were vortexed for 15 s and submitted to cell disruption for 15-min at 4°C using a TissueLyser II (Qiagen) at 30 Hz. Tubes were then centrifuged for 5 min at 4°C. Resulting supernatants were transferred to new tubes, and 200 μL of chloroform was added. After vortexing for 15 s, the solution was for 5 min at room temperature. RNA extraction was conducted in accordance with manufacturer’s guidelines. Resulting extracts were resuspended in 50 μL RNAse-free water and underwent DNA removed using the Turbo DNase kit (Invitrogen). Lastly, samples were concentrated through ethanol precipitation following previously outlined protocol (Green and Sambrook, 2016), and resuspended in RNAse-free water to a final volume of 30 μL. Sample concentration and purity were monitored by NanoDrop 2000 (Thermo Fisher Scientific) at all steps. Quality of extracted RNAwas assessed by electrophoresis on 1.5% (w/v) agarose gels. Prior to library preparation, the quality of the extracts was verified using the 2100 Bioanalyzer (Agilent) and NanoDrop 2000. Subsequently, PolyA enrichment, cDNA library construction, paired-end sequencing using an Illumina NovaSeq with 150 bp read length, quality control, and sequence trimming were conducted by NovogeneAIT.

### 2.6 Transcriptomic analysis

Trimmed reads were aligned to *R. toruloides* NRRL Y-1091 genome (NCBI GenBank accession number: GCA_001542305.1) using STAR v2.7.7 (Dobin *et al*., 2013). Subsequently, RSEM v1.3.3 (Li and Dewey, 2011) was used to quantify gene expression levels with reference to genome annotation generated by AUGUSTUS v3.3.3 (Stanke and Morgenstern, 2005). Expression levels are quantified in counts of Fragments Per Kilobase of transcript per Million mapped reads (FPKM) or Transcripts Per Million (TPM). Counts processing and differential expression analysis (DE) were performed using DEseq2 v1.30.0 (Love *et al*., 2014). Initially, data filtering was done by retaining genes with TPM above 10 in at least three samples. Filtered data was normalized to adjust for variations in sequencing depth through variance stabilizing transformation (Anders and Huber, 2010). Hierarchical clustering and principal component analysis (PCA) utilized the normalized data, whereas TPM values were employed for DE. Significantly differentially expressed genes were defined as genes with an adjusted p-value less than 0.01. In the case of PCA and hierarchical clustering, biological replicates were not combined. Instead, the z-scores of normalized counts for pertinent genes across various conditions were utilized to assess inter-replicate variability.

### 2.7 Genomic analysis

We identified genes associated with glycerol metabolism, transport, and lipid biosynthesis through functional annotations, KO numbers (Mao *et al*., 2005) and by searching for homologous genes in other yeast species using the AYbRAH database (Correia *et al*., 2019). To identify common responsive elements in the promoters of relevant genes, 1000 bp upstream and downstream of coding sequences were extracted using the FlankBed and GetFastaBed (Bedtools) (Quinlan and Hall, 2010) from GALAXY server (Giardine *et al*., 2012). Promoter sequences (upstream or downstream) were selected based on the gene orientation and compared in search for common motifs using info-Gibbs (Defrance and van Helden, 2009). Identified motifs were compared to *S. cerevisiae* as well as fungal responsive element and cross-checked using compare-matrices and matrix-scan from the Regulatory Sequence Analysis Tools, respectively (Nguyen *et al*., 2018).

## 3. Results and Discussion

### 3.1 Investigating the glycerol tolerance of oleaginous yeasts

21 strains of 17 yeast species (**Table 1**) which have high industrial potential for lipid production were selected. Firstly, microbiology-grade glycerol which will have no interference from impurities present in sweetwater was used to determine strain tolerance to high glycerol concentrations. The strains were cultivated in media containing different concentrations of glycerol (2%, 8%, and 16% w/v), supplemented with nutrients using SC mix. These media were denoted as Gly_2_SC, Gly_8_SC, and Gly_16_SC, respectively, to reflect their glycerol concentration and composition (**Table 2** and **Fig. S2**). The growth curves obtained were analyzed to derive carrying capacities (CC) and t_mid_, which represents the maximum population size attained by the available carbon sources and the time required to reach half of the CC, respectively.

**Table 2.**
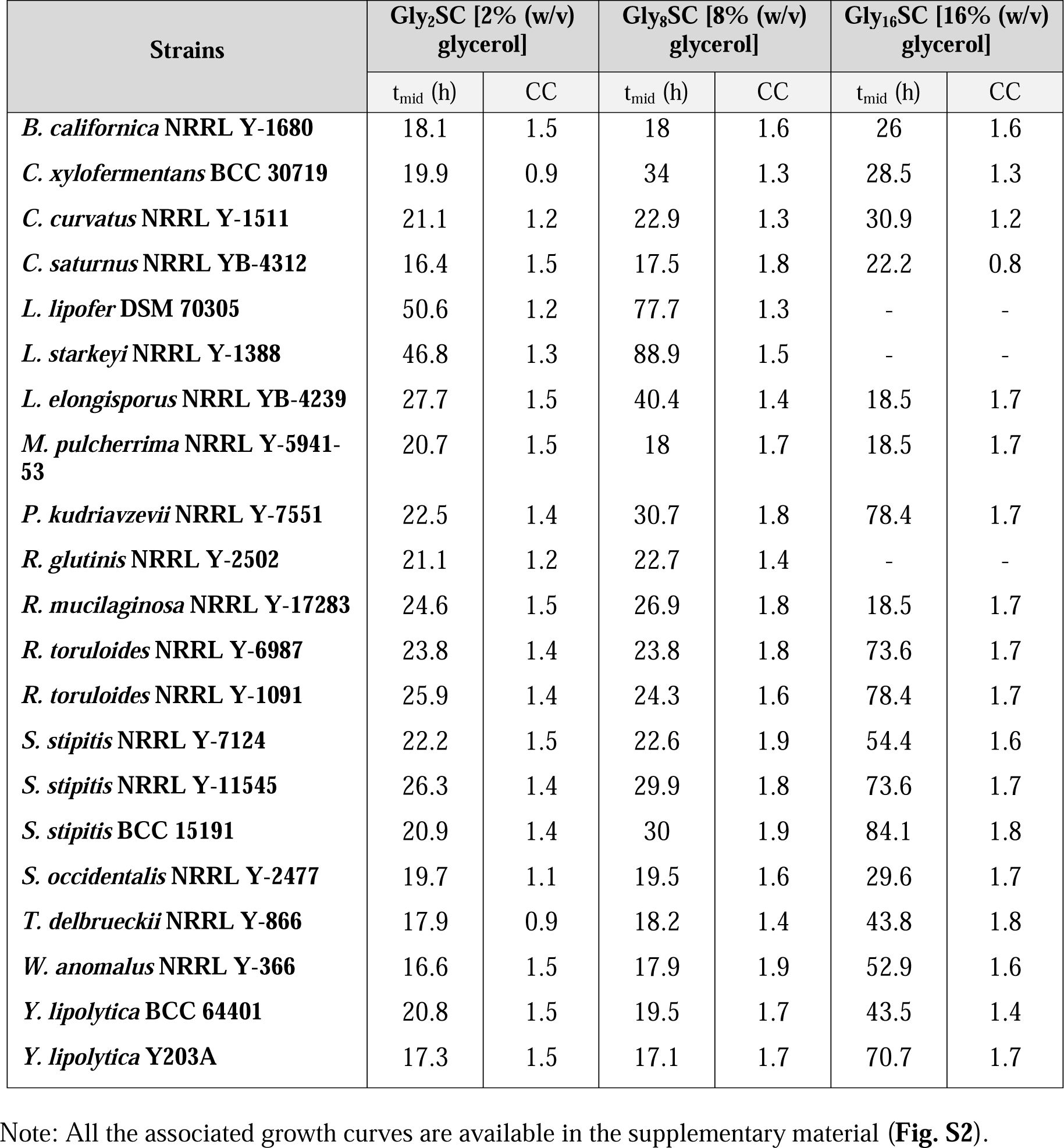
Strains tolerance to microbiology-grade glycerol.

| Strains | Gly<sub>2</sub>SC [2% (w/v) glycerol] |  | Gly<sub>8</sub>SC [8% (w/v) glycerol] |  | Gly<sub>16</sub>SC [16% (w/v) glycerol] |  |
| --- | --- | --- | --- | --- | --- | --- |
|  | t<sub>mid</sub> (h) | CC | t<sub>mid</sub> (h) | CC | t<sub>mid</sub> (h) | CC |
| *B. californica* NRRL Y-1680 | 18.1 | 1.5 | 18 | 1.6 | 26 | 1.6 |
| *C. xylofermentans* BCC 30719 | 19.9 | 0.9 | 34 | 1.3 | 28.5 | 1.3 |
| *C. curvatus* NRRL Y-1511 | 21.1 | 1.2 | 22.9 | 1.3 | 30.9 | 1.2 |
| *C. saturnus* NRRL YB-4312 | 16.4 | 1.5 | 17.5 | 1.8 | 22.2 | 0.8 |
| *L. lipofer* DSM 70305 | 50.6 | 1.2 | 77.7 | 1.3 | - | - |
| *L. starkeyi* NRRL Y-1388 | 46.8 | 1.3 | 88.9 | 1.5 | - | - |
| *L. elongisporus* NRRL YB-4239 | 27.7 | 1.5 | 40.4 | 1.4 | 18.5 | 1.7 |
| *M. pulcherrima* NRRL Y-5941-53 | 20.7 | 1.5 | 18 | 1.7 | 18.5 | 1.7 |
| *P. kudriavzevii* NRRL Y-7551 | 22.5 | 1.4 | 30.7 | 1.8 | 78.4 | 1.7 |
| *R. glutinis* NRRL Y-2502 | 21.1 | 1.2 | 22.7 | 1.4 | - | - |
| *R. mucilaginosa* NRRL Y-17283 | 24.6 | 1.5 | 26.9 | 1.8 | 18.5 | 1.7 |
| *R. toruloides* NRRL Y-6987 | 23.8 | 1.4 | 23.8 | 1.8 | 73.6 | 1.7 |
| *R. toruloides* NRRL Y-1091 | 25.9 | 1.4 | 24.3 | 1.6 | 78.4 | 1.7 |
| *S. stipitis* NRRL Y-7124 | 22.2 | 1.5 | 22.6 | 1.9 | 54.4 | 1.6 |
| *S. stipitis* NRRL Y-11545 | 26.3 | 1.4 | 29.9 | 1.8 | 73.6 | 1.7 |
| *S. stipitis* BCC 15191 | 20.9 | 1.4 | 30 | 1.9 | 84.1 | 1.8 |
| *S. occidentalis* NRRL Y-2477 | 19.7 | 1.1 | 19.5 | 1.6 | 29.6 | 1.7 |
| *T. delbrueckii* NRRL Y-866 | 17.9 | 0.9 | 18.2 | 1.4 | 43.8 | 1.8 |
| *W. anomalus* NRRL Y-366 | 16.6 | 1.5 | 17.9 | 1.9 | 52.9 | 1.6 |
| *Y. lipolytica* BCC 64401 | 20.8 | 1.5 | 19.5 | 1.7 | 43.5 | 1.4 |
| *Y. lipolytica* Y203A | 17.3 | 1.5 | 17.1 | 1.7 | 70.7 | 1.7 |

Note: All the associated growth curves are available in the supplementary material (**Fig. S2**).

All yeast strains demonstrated the ability to utilize Gly_2_SC, and some were capable of tolerating Gly_8_SC. *T. delbrueckii*, *S. occidentalis*, *W. anomalus*, *R. toluroides* NRRL Y-6987, *S. stipilis* NRRL Y-11545, *S. stipilis* BCC 15191 and *C. xylofermentans* exhibited improved performance in Gly_8_SC, achieving a higher final OD_600_ while maintaining a similar growth rate to that observed in Gly_2_SC, suggesting that these strains are less susceptible to substrate inhibition. *C. xylofermentans*, *S. occidentalis*, and *T. delbruekii* demonstrated a CC in Gly_8_SC 1.5 times higher than that observed in Gly_2_SC. However, all strains were unable to achieve higher cell densities in Gly_16_SC, indicating that they were inhibited at high glycerol concentration (**Table 2**). Despite this, *L. elongisporus*, *M. pulcherrima*, and *R. mucilaginosa* are the most tolerant strains, achieving a similar t_mid_ value for both Gly_2_SC and Gly_16_SC. In addition, *T. delbrueckii*, *S. occidentalis* and *R. toluroides* NRRL Y-6987 were also able to produce a final biomass which is comparable to Gly_2_SC and Gly_8_SC.

### 3.2 Evaluating yeast ability to utilize industrial sweetwater

Next, yeasts were evaluated based on their abilities to utilize sweetwater for growth. Three sweetwater types are available: SW_15_, SW_35_ and SW_85_, which were collected from several points along the glycerol refining chain. They have increasing glycerol concentration and purity. Previously, dilution of glycerol-containing discharge from biodiesel manufacturing process with synthetic media was shown to improve microbial growth (Ito *et al*., 2005). However, this is impractical at industrial scale due to the associated increase in production cost. Hence, the use of SW_15_, the crudest form of sweetwater with a relatively lower concentration of glycerol without prior dilution is preferable in terms of cost. Conversely, although SW_35_ and SW_85_ offer the advantages such as higher purity and lower moisture levels, rendering them an extended shelf life, dilution and/or nutrient addition prior use is necessary due to their higher glycerol concentration.

SW_15_, SW_35_ and SW_85_ were supplemented with SC mix to produce media with a final glycerol content of 1.5 % (w/v), referred to as SW_15>1.5_SC, SW_35>1.5_SC and SW_85>1.5_SC, respectively. We first compared the suitability of these three media for yeast cultivation. For most yeast strains, growth profiles were similar regardless of how the diluted sweetwater was prepared. Additionally, the growth profiles of all yeast strains in these media were highly similar to that of Gly_2_SC (**Table 2**). For simplicity, SW_15>1.5_SC, SW_35>1.5_SC and SW_85>1.5_SC were collectively referred to as SW_1.5_SC, and this medium served as a reference medium throughout this study. Next, we evaluated the growth of all 21 strains in SW_15_ to identify differences resulting from sweetwater dilution and/or nutrient supplementation. To our surprise, all 21 strains were able to grow in SW_15_ despite its high glycerol concentration, including *L. lipofer* and *L. starkeyi* which were unable to grow in Gly_16_SC (**Table 3**), potentially suggesting the presence of other preferred carbon sources in sweetwater. However, the OD_600_ recorded in SW_15_ was slightly lower than SW_1.5_SC. This observation further validated growth inhibition at high glycerol concentration. Nevertheless, we can confirm that SW_15_ is suitable for yeast growth without nutrient supplementation. In fact, the composition of SW_15_ enhanced the growth of some yeast strains. *R. glutinis* showed low slow growth in Gly_16_SC but achieved one of the highest biomass in SW_15_. *R. mucilaginosa* and *R. toruloides* NRRL Y-6987 growth rates were also improved in SW_15_, but their carrying capacities decrease, indicating that SW_15_ initially offered an environment more favorable for proliferation, but later turned inhibitory possibly due to low initial pH of SW_15_ (pH 3.95) or depletion of nutrients.

**Table 3.** Effects of glycerol concentration and pH of sweetwater on yeast growth.

| Strains | Concentration effect |  |  |  | pH effect |  |  |  |
| --- | --- | --- | --- | --- | --- | --- | --- | --- |
|  | SW<sub>35>1.5</sub>SC |  | SW<sub>35</sub>SC |  | SW<sub>15</sub><br>pH 6.09 |  | SW<sub>15</sub><br>pH 3.95 |  |
|  | t<sub>mid</sub> (h) | CC | t<sub>mid</sub> (h) | CC | t<sub>mid</sub> (h) | CC | t<sub>mid</sub> (h) | CC |
| *B. californica* NRRL Y-1680 | 9.4* | 1.4* | 19.7 | 1.6 | 21.2 | 1.05 | 22.7 | 1.05 |
| *C. xylofermentans* BCC 30719 | 23.7 | 1 | 60.8 | 1.9 | 39.7 | 1.2 | 49.2 | 0.99 |
| *C. curvatus* NRRL Y-1511 | 21.3 | 1.5 | 65 | 1.4 | 41.5 | 1.24 | 93.6 | 1.29 |
| *C. saturnus* NRRL YB-4312 | 13.4 | 1.5 | 26.1 | 1.7 | 29.2 | 1.32 | 31.3 | 1.29 |
| *L. lipofer* DSM 70305 | 59.5 | 0.9 | 171.9 | 0.7 | 79.7 | 0.8 | 66.1 | 0.35 |
| *L. starkeyi* NRRL Y-1388 | 47.1 | 1 | - | - | 66.8 | 0.75 | 67.9 | 0.59 |
| *L. elongisporus* NRRL YB-4239 | 18.3 | 1.3 | 35.9 | 1.8 | 21.5 | 1.08 | 19.9 | 1.15 |
| *M. pulcherrima* NRRL Y-5941-53 | 19.1 | 1.4 | 38.3 | 1.7 | 19.8 | 1.41 | 20.2 | 1.42 |
| *P. kudriavzevii* NRRL Y-7551 | 22.1 | 1.3 | 37 | 1.5 | 27.6 | 0.87 | 54.1 | 0.73 |
| *R. glutinis* NRRL Y-2502 | 24.2 | 1.3 | 103.3 | 1.5 | 27.2 | 1.29 | 54.2 | 1.32 |
| *R. mucilaginosus* NRRL Y-17283 | 20 | 1.3 | 22.4 | 1.7 | 20.6 | 1.1 | 23.8 | 1.02 |
| *R. toruloides* NRRL Y-6987 | 23 | 1.6 | 35.4 | 1.7 | 21.7 | 1.34 | 28.1* | 1.28* |
| *R. toruloides* NRRL Y-1091 | 21.5 | 1.4 | 55.8 | 1.8 | 24.1 | 1.15 | 42.4 | 1.13 |
| *S. stipitis* NRRL Y-7124 | 20.1 | 1.4 | 139.2 | 1.8 | 28.1 | 1.3 | 21.7 | 1.08 |
| *S. stipitis* NRRL Y-11545 | 19.4 | 1.4 | 37 | 1.8 | 21.3 | 1.23 | 23.9 | 1.09 |
| *S. stipitis* BCC 15191 | 23.8 | 1.3 | 33.7 | 1.7 | 20.7 | 1.26 | 21.2 | 1.17 |
| *S. occidentalis* NRRL Y-2477 | 19.4 | 1.1 | 24.2 | 1.7 | 33.8 | 1.45 | 49 | 1.24 |
| *T. delbrueckii* NRRL Y-866 | 18 | 0.8 | 26.6 | 1.6 | 21.6 | 1.02 | 21.1 | 0.57 |
| *W. anomalus* NRRL Y-366 | 13.7 | 1.4 | 122.5 | 2 | 20.3 | 1.41 | 21.7 | 1.47 |
| *Y. lipolytica* BCC 64401 | 20.4* | 1.4* | 20 | 1.8 | 21.6 | 1.21 | 20.7 | 1.32 |
| *Y. lipolytica* Y203A | 17.3* | 1.5* | 18.3 | 1.7 | 32.8 | 1.33 | 19.8 | 1.32 |

Note: All the associated growth curves are available in the supplementary material (**Fig. S3 and** **S4**). An asterisk (\*) indicates averages computed with less than 4 replicates.

To further explore optimal growth conditions, we formulated an additional media using SW_15_ by adjusting its pH from 3.95 to 6.09. Previous research suggested that pH adjustments can affect yeast growth in glycerol (Chiruvolu *et al*., 1998; Swinnen *et al*., 2013). However, we found that the average growth rate of the strains was not significantly different at both pH values, although certain individual differences were observed. Specifically, *R. glutinis* NRRL Y-2502, *R. toruloides* NRRL Y-1091, *C. curvatus*, *T. delbrueckii*, and *S. occidentalis* performed better in pH 6.09, while *Y. lipolytica* (20, 21) strains slightly favored low pH. Overall, in both pH conditions, *W. anomalus*, *M. pulcherrima*, and *Y. lipolytica* (20, 21) exhibited the highest carrying capacities and fastest growth rates, indicating that they are among the best yeast candidates for sweetwater utilization. Since growth differences in two pH conditions were minor, we can infer that nutrient limitation is the limiting factor for yeast growth in SW_15_.

### 3.3 Assessing lipid production of 21 strains grown in SW_15_ and SW_1.5_SC

We next compared the lipid production of 21 strains in SW_15_. To address the nutrient limitation in SW_15_, we also included SW_1.5_SC. Furthermore, taking into consideration the varying growth rates of different strains and the coupling of lipid accumulation to growth phase, we analyzed the cellular lipid content after 3 and 10 days of cultivation. Based on average fluorescence signals across the 4 samples (SW_15_ 3 days, SW_15_ 10 days, SW_1.5_SC 3 days, SW_1.5_SC 10 days), we found that the best lipid accumulators are *R. toluroides* NRRL Y-6987, *L. starkeyi*, *R. glutinis*, *C. saturnus* and *L. elongisporus* (**Fig. 1a**). Notably, some strains had higher lipid accumulation in SW_15_. For instance, *R. mucilaginosa*, *S. stiptitis* (14), *M. pulcherrima* and *C. saturnus* accumulated more lipids in SW_15_ in comparison to SW1.5SC on average (**Fig. 1a**), suggesting that these strains better accumulate lipids under low nutrient conditions and/or high C/N ratio. Important to highlight, nutrient supplementation will also likely reduce the C/N ratio. On the other hand, *W. anomalus*, *Y. lipolytica* (20, 21) and *M. pulcherrima* showed relatively low lipid accumulation, despite growing best in sweetwater.

**Fig. 1.**
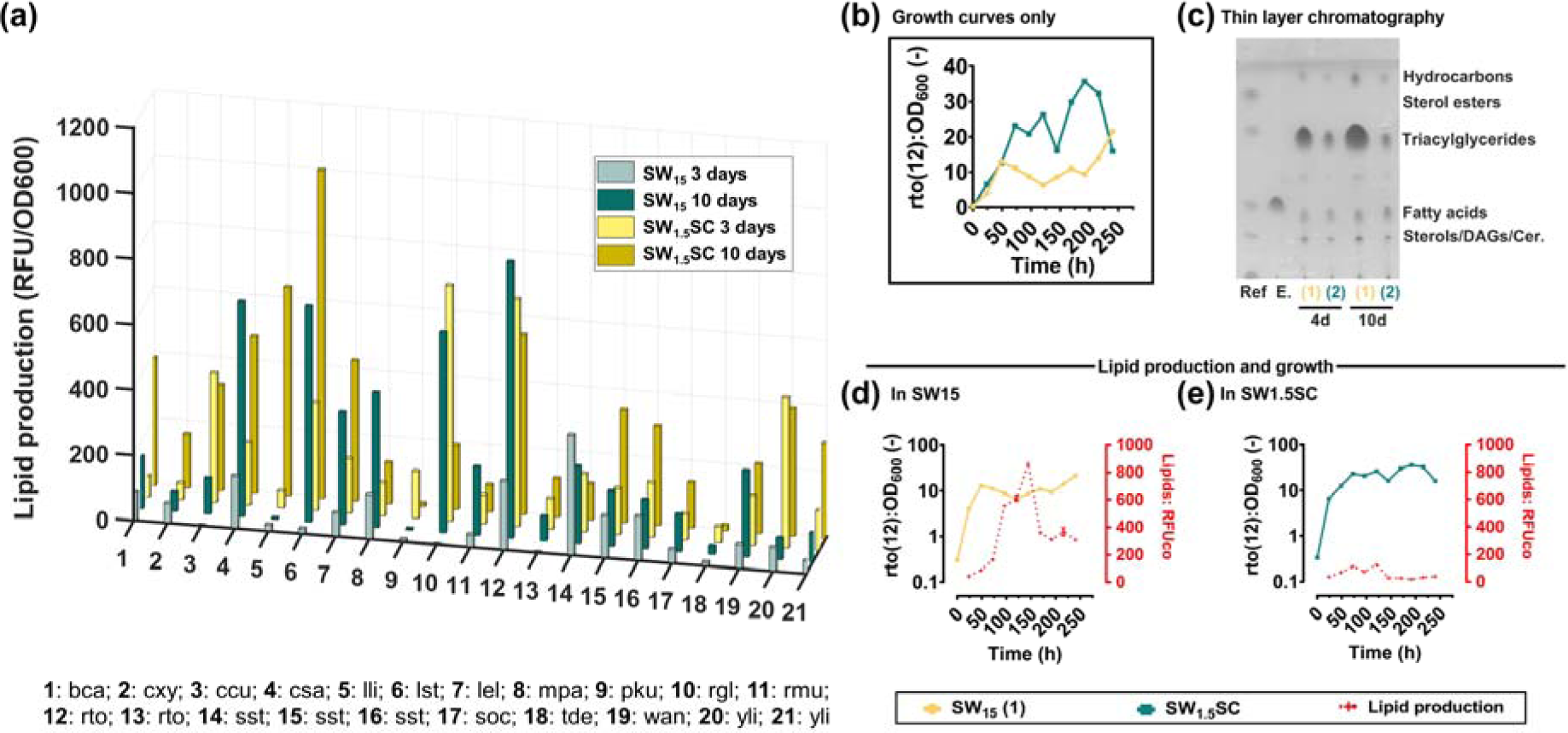
Lipid production in sweetwater-based media. (a) Comparison of lipid production in SW_15 s_the 21 investigated strains. 3-letter codes are abbreviations for full species names which can be found in **Table 1**. For clarity, standard deviation was not shown in the bar chart. (b) Comparison of *R. toluroides* NRRL Y-6987 growth in different media. (c) Comparison of *R. toluroides* NRRL Y-6987 lipid accumulation profiles in different media. (d-e) Lipid content over time. OD_600_ were plotted on a logarithmic scale for improved visualization. Fluorescence of the stained cells was measured in quadruplicate and subsequently normalized by the OD600 of each sample, resulting in RFU/OD600 values.

Since *W. anomalus* and *R. toluroides* are the strains with the highest growth and lipid accumulation in sweetwater-based medium, respectively, we proceeded to verify their growth and SW-induced lipid accumulation abilities through time-course monitoring at a larger scale cultivation. Shake-flask cultivation confirmed that *W. anomalus* achieves higher OD_600_ (data not shown) than *R. toluroides* NRRL Y-6987 when grown in the same conditions. However, lipid accumulation in SW_15_ of *R. toluroides* NRRL Y-6987 is 3.9 times higher than *W. anomalus*. Furthermore, maximum normalized fluorescence signals recorded for *R. toluroides* NRRL Y-6987 in SW_15_ is 7.1× higher than that in SW_1.5_SC, demonstrating that lipid accumulation of *R. toluroides* NRRL Y-6987 is better induced by lower nutrient conditions and/or higher C/N ratio in SW_15_ (**Fig. 1d-e**). Such lipid accumulation difference is also clearly visible on TLC (**Fig. 1c**). Taken together, this indicates that nutrient supplementation or dilution is not necessary for lipid accumulation. From industrial application perspective, SW_15_ can be directly used for microbial fermentation for lipid production without pre-processing.

Growth in SW_15_ (**Table 3**) suggests the presence of nitrogen sources, which give a favorable C:N ratio for lipid accumulation for some strains. Nitrogen limitation is reported to reduce the intracellular levels of adenosine monophosphate (AMP) (Yoshino and Murakami, 1982), thereby inhibiting isocitrate dehydrogenase in the Krebs cycle which induces accumulation of citrate in mitochondria, which is then exported to the cytoplasm for acetyl-CoA production through citrate lyase (ACL) (Adrio, 2017). In some yeasts, citrate accumulation is known to initiate fatty acid synthesis by activation of acetyl-CoA carboxylase (Botham and Ratledge, 1979).

Furthermore, impurities in sweetwater including free fatty acids, monoacylglycerol, diacylglycerol and triacylglycerol can function as a secondary carbon source or add to the intracellular lipid pool, thereby contributing to growth or lipid accumulation. In fact, *C. curvatus*, *R. toruloides* and *Y. lipolytica* (Gao *et al*., 2016; Matatkova *et al*., 2017; Patel and Matsakas, 2019) was reported to utilize hydrophobic substrates. Taken together, since SW_15_ can promote lipid accumulation, approaches such as optimizing feeding strategies or improving strain tolerance to high glycerol concentration by adaptive laboratory evolution can be applied to improve cell densities in SW_15_ (González-Villanueva *et al*., 2019).

### 3.4 Investigating the transcriptomic changes related to sweetwater utilization by *R.* toruloides (NRRL Y-6987)

To elucidate the mechanism underlying sweetwater-induced lipid accumulation, we formulated four different media using sweetwater or microbiology-grade glycerol, each with or without nutrients, respectively named SW_15_, SW_1.5_SC, Gly_15_ and Gly_1.5_SC. Next, RNA from *R. toruloides* (NRRL Y-6987) grown in these four media was extracted and sequenced, followed by analysis of the transcriptomic changes (see **Material and Methods**). PCA showed clear separation between the conditions and the close clustering of the biological replicates of each condition. PCA also confirmed that the predominant factor driving the observed differences was nutrient availability, as the first principal component (PC1) accounted for 60.4% of the total variation observed and clearly separated samples in media with or without supplementation of nutrients (**Fig. 2a**). Next, the second principal component (PC2) which explains 17.4% of the variations clearly separated the transcriptomes of samples from sweetwater and microbiology-grade glycerol. This is observed from projection on PC2 which produced the greatest distance between Gly_1.5_SC and SW_1.5_SC. The third principal component (PC3) which accounts for 11.5% of the differences also represents the differences between microbiology-grade glycerol and sweetwater (**Fig. 2b**), evidenced by SW_15_ and Gly_15_ transcriptomes which clustered furthest away from each other. These results indicate that supplementation of nutrients, followed by the source of glycerol, are key factors for transcriptomic changes.

**Fig. 2.**
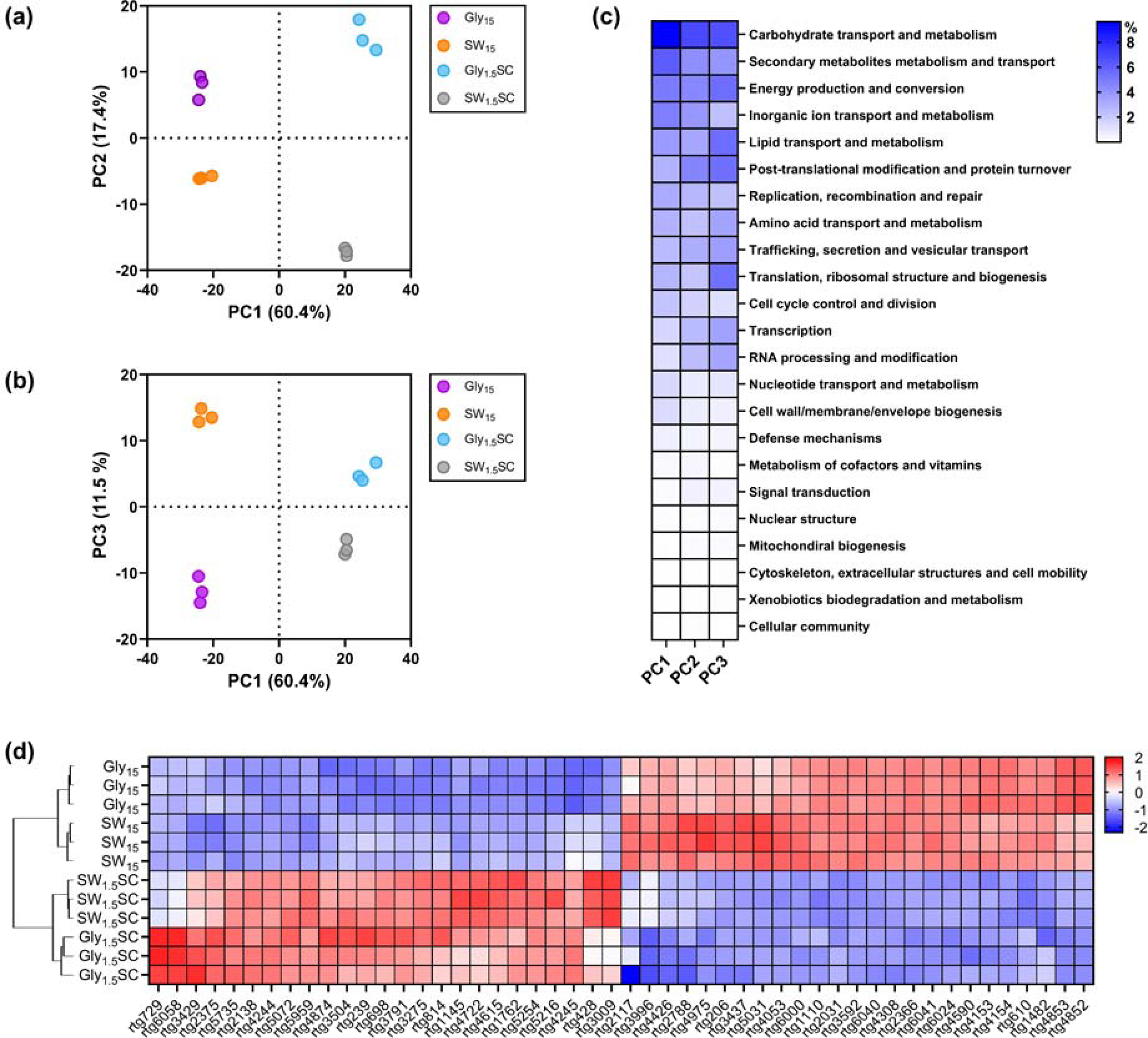
Comparison of *R. toruloides* (NRRL Y-6987) transcriptome in 4 glycerol-based media. (a-b) PCA score plots of transcriptomes in Gly_15_, SW_15_, Gly_1.5_SC and SW_1.5_SC. (c) Functional categories were sorted by descending contribution. To improve visibility, genes with unknown function were excluded from the heatmap. (d) Heatmap showing normalized expression counts of the top 50 annotated genes which contributes the most to PC1-3.

Next, all genes were categorized into 24 functional groups and their contributions to each category to PC1, PC2 and PC3 was computed (**Fig. 2c**). We found that genes contributing the most to PC1, PC2 and PC3 are in the categories of “Unknown function”, “Carbohydrate transport and metabolism”, “Secondary metabolites metabolism and transport”, “Energy production and conversion”, “Inorganic ion transport and metabolism” and “Lipid transport and metabolism”. Predominance of genes of unknown function highlights the importance of characterizing unconventional yeasts, as we did in this study. Even so, we can infer that metabolism and transport of carbohydrates, secondary metabolites and inorganic ions are most affected by nutrient availability. Important to note, SC nutrient mix enriches the growth media with essential amino acids, nitrogenous bases, inorganic ions, and vitamins, and therefore may cause gene expression changes in the relevant gene categories. For example, inorganic ion transport and metabolism which contributed significantly to PC1. On the other hand, genes involved in energy and lipid metabolism contribute more to PC3, suggesting major differences in related pathways when cells are grown in sweetwater as compared to microbiology-grade glycerol. Next, “Lipid metabolism and transport” contains genes involved in lipid biosynthesis and degradation, as well as lipid modification and mobilization. Additional observations on the globally low contribution of the category “Cell wall/membrane/envelope biogenesis” suggests that the high contribution of lipid metabolism is mainly due to their involvement in signaling processes and lipid storage instead of involvement in membrane biogenesis/modification.

Hierarchical clustering confirmed that *R. toluroides* cultivated in sweetwater and glycerol media exhibited different transcriptomic responses (**Fig. 2d** and **Fig. S5**). To identify the genes behind the transcriptomic changes associated with nutrient availability and glycerol type, we analyzed the top 50 functionally annotated genes contributing the most to PC1, PC2 and PC3. Hierarchical clustering was performed on the z-scores of normalized gene expression of the top 50 genes to visualize genes with high inter-replicate variability (**Fig. 2d**). We found that, for a given condition, gene expression across replicates was highly consistent, which is coherent with the close clustering among biological replicates observed in PCA, validating the quality and consistency of our data. Next, we compared expression levels of the top 50 genes to investigate the variances between two experimental conditions: (1) absence of nutrient addition and high glycerol concentration (SW_15_ and Gly_15_) and (2) nutrient addition with low glycerol concentration (SW_1.5_SC and Gly_1.5_SC).

We observed that genes involved in carbohydrate transport and metabolism, namely *rtg3791* (chitinase), *rtg1145* (TNA1), *rtg1762* (HXT), *rtg5254* (HXT), *rtg4244* (MAL31), *rtg4245* (IMA1), *rtg4615* (ecfuP), and *rtg4722* (JEN1) are more expressed in the presence of nutrients at low glycerol concentrations (**Fig. 2d**). Since the cell wall of *R. toruloides* contains chitin (Buck and Andrews, 1999), *rtg3791* likely contributes to cell wall remodeling activities. Remaining genes can be identified as proton symporters for hexoses (*rtg1762* and *rtg5254*), maltose (*rtg4244*), fucose (*rtg4615*) and carboxylic acids (*rtg4722*) (**Table S1**), while JEN1 is a lactate transporter induced by non-fermentable carbon sources including glycerol, derepressed by glucose absence (Lodi *et al*., 2004). Interestingly, expression of JEN1 is dependent on the kinase SNF1 (Chambers *et al*., 2004), which connects various pathways involved in stress regulation including TORC pathway for nitrogen sensing, RTG2/SNF3 pathway for glucose sensing, PKA pathway for general stress response and HOG1 pathway for osmotic shock. Next, most lipid metabolism genes with high contribution to PC1 to PC3 are putatively involved in lipid degradation and modification, such as acyl-CoA dehydrogenase (*rtg5959*), peroxisomal MaoC dehydratase (*rtg2138*), and peroxisomal 2,4-dienoyl-CoA reductase (*rtg206*). Since acyl-CoA dehydrogenase is involved in β-oxidation, this may suggest that β-oxidation is more active in the presence of nutrients at low glycerol concentration (**Table S1**).

Comparing Gly_15_ and SW_15_ which both have no nutrient addition and high glycerol concentration, genes related to ion transport highly contributes to the PCs, in particular *rtg6000* (ammonium transporter, Amt family) which is involved in NH_4_^+^/NH_3_ uptake (**Fig. 2d**; **Table S1**). Notably, *rtg6000* which was upregulated in SW_15_ is homologous to three endogenous scMEP proteins of *S. cerevisiae* and shares the highest identity with scMEP2, an ammonium sensor which induces pseudohyphal growth during ammonium limitation (Boeckstaens *et al*., 2007). scMEP genes encode ammonia permeases and are highly upregulated after exhaustion of a preferred nitrogen source or in presence of non-preferred source (Crépin *et al*., 2014). These genes are also related to nitrogen catabolite repression which allows selection of the optimal nitrogen source for growth (Beltran *et al*., 2004). This suggests that SW_15_ and Gly_15_ differ in their nitrogen composition. Taken together, *R. toruloides* (NRRL Y-6987) responses to the change of growth environment by adjusting primarily its carbohydrate, inorganic ion, and lipid metabolism.

### 3.5 Comparative analysis for SW_15_ and Gly_15_

To understand more about the transcriptomic differences between SW_15_ and Gly_15_ and investigate possible mechanisms underlying SW-induced lipid accumulation, a second analysis was performed by focusing only on the relevant transcriptomes. Growth is improved substantially in SW_15_ in comparison to Gly_15_ **(Fig. S6)**. 72.92% of the difference between the two conditions can be captured by PC1 while other PCs mainly describe inter-replicate differences (**Fig. 3b**). The top 50 annotated genes contributing the most to PC1 was also comprehensively analyzed.

**Fig. 3.**
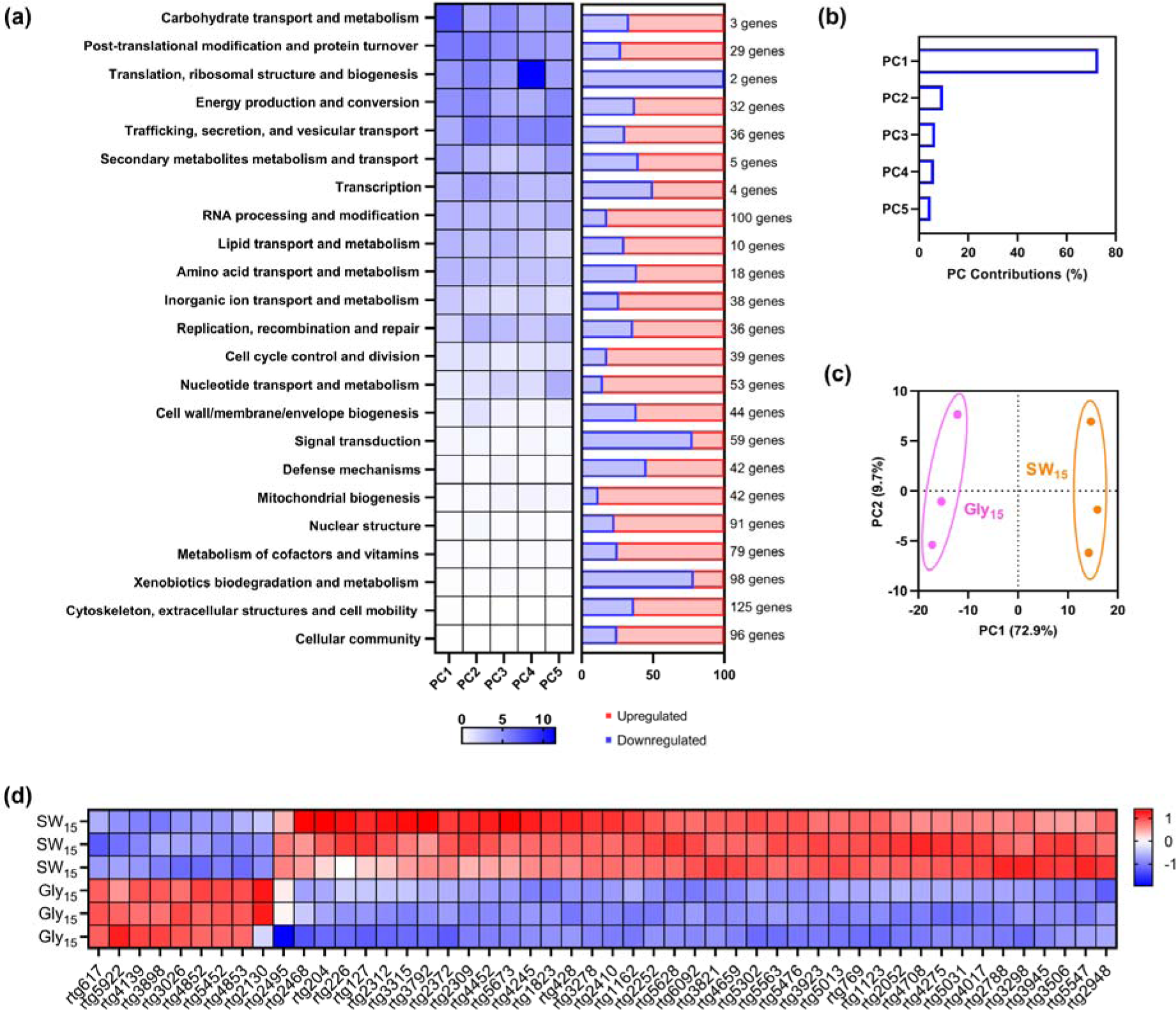
Comparison of R. toruloides (NRRL Y-6987) transcriptome in SW15 and Gly15. (a) Heatmap on the left shows the contribution of each functional category to the principal components (PC1-5). The functional categories were sorted in a descending order of global contribution to the PCs. In the right part, the proportion of up- or down-regulated genes compared to Gly_15_ is given for each functional category. (b) Scree plot shows the percentage of variance explained by each PC. (c) PCA score plots of transcriptomes in Gly15 and SW15 PCA score plots. (d) Normalized expression counts of the top 50 annotated genes contributing the most to PC1. The heatmap represents row-standardized z-scores.

Our data and analysis again hinted the presence of nitrogen source(s) in SW_15_. *rtg4852* and *rtg4853*, respectively annotated as nitrate/nitrite transporter and nitrite reductase were highly expressed in Gly_15_ (**Fig. 3a** and **Fig. 4**). *rtg4853* shares homology with umNAR1 from *Ustilago maydis* and ncNIT-6 from *Neurospora crassa*, genes which are known to be influenced by nitrogen metabolite repression, and are triggered by the absence of ammonia or presence of nitrate (Banks *et al*., 1993; Johns *et al*., 2016). On the other hand, in SW_15_, the low expression of *rtg4853* suggests the presence of ammonium or nitrogen source in SW_15_. Oligopeptides may be a possible nitrogen source, indicated by increased expression of *rtg3315* and *rtg5673* which are oligopeptide transporters. Furthermore, upregulation of *rtg6000* which encodes an ammonium transporter and the amidase *rtg3792* which facilitates ammonium production from arginine, tryptophan and phenylalanine implies the digestion of exogenous oligopeptides and subsequent processing of resulting amino acids (**Fig. 4** and **Table S2**). Additionally, *rtg5216* (ubiquitin C) expressing 5-times more in SW_15_ further supports that *rtg3792* is metabolizing amino acids, leading to differences in protein turnover/autophagy.

**Fig. 4.**
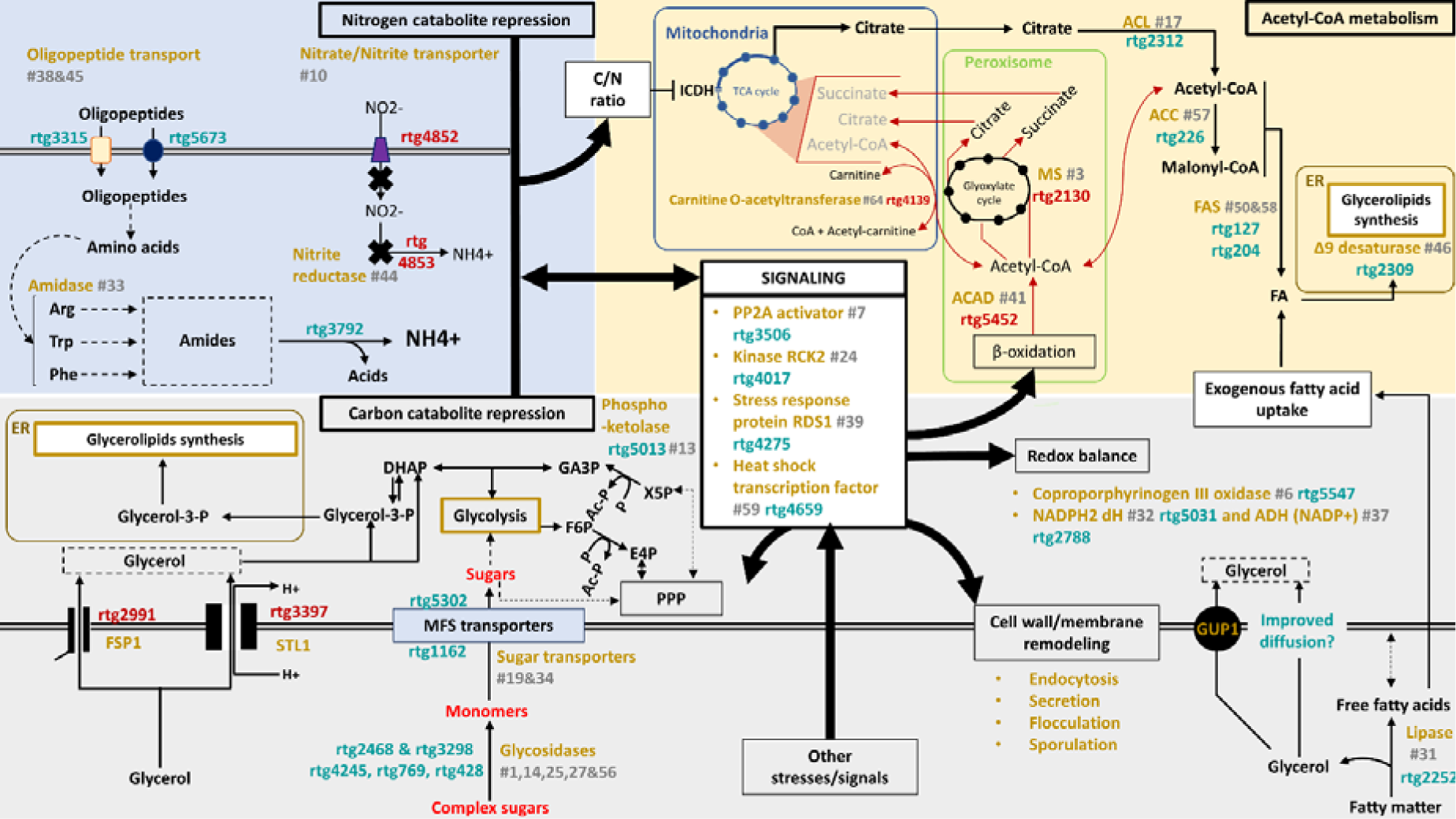
Scheme summarizing the main differences between the growths of *R. toruloides* (NRRL Y-6987) in SW_15_ and in Gly_15_. Genes upregulated in SW_15_ (Gly_15_ as control) are annotated in teal while genes downregulated in SW_15_ (Gly_15_ as control) are annotated in crimson. The function or enzyme names are shown in yellow, followed by its rank in terms of contribution to PC1, shown in grey. A dashed arrow represents a potential relation, while solid lines signify a high level of confidence. <u>Abbreviations:</u> ACAD, acyl-CoA dehydrogenase; ACC, acetyl-CoA carboxylase; ACL, ATP citrate lyase; ADH, alcohol dehydrogenase; Ac-P, acetyl-phosphate; dH, dehydrogenase; DHAP, dihydroxyacetone; ER, endoplasmic reticulum; E4P, Erythrose-4-phosphate; FAS, Fatty acid synthase; F6P, Fructose-6-phosphate; GA3P, Glyceraldehyde-3-phosphate; ICDH, isocitrate dehydrogenase; MFS, Major facilitator superfamily; MS, malate synthase; P, phosphate; PPP, Pentose phosphate pathway; PP2A, protein phosphatase 2A; RCK, Radiation sensitivity Complementing Kinase; RDS, Regulator of Drug Sensitivity; X5P, Xylulose-5-Phosphate.

Conversely, when nitrogen is limited, citrate accumulates as a result of a less active Krebs cycle. Excess citrate is then directed into fatty acid synthesis. Later, acetyl-CoA is produced from citrate through ACL, then converted into malonyl-CoA by acetyl-CoA carboxylase (ACC). These two enzymes, along with 2 fatty acid synthases subunits (α: *rtg127*; β: *rtg204*) **(Fig. 4)** and 9 other enzymes related to glycerolipids, phospholipids and carotenoid production (**Table S3**) are upregulated in sweetwater. This is consistent with a higher lipid accumulation in SW_15_, which is not observed in Gly_15_. This may be explained by higher expression of β-oxidation in Gly_15_ than in SW_15_. This process can release peroxisomal acetyl-CoA, which in turn supplies the glyoxylate cycle and potentially counteracts the ICDH-induced decline in the Krebs cycle, thus blocking the re-direction of citrate towards lipid production (**Fig. 4**). Therefore, cells likely produce energy by redirecting acetyl-CoA flux towards TCA cycle in Gly_15_. In SW_15_, reduced expression of β-oxidation related proteins such as acyl-CoA dehydrogenase (*rtg5452*), carnitine o-acyltransferase (*rtg4139*), and malate synthase (*rtg2130*), a key enzyme of the glyoxylate cycle, supports our hypothesis.

Other than the presence of nitrogen source, SW_15_ also includes some tri-/di-/mono-glycerides and free fatty acids, which are by-products of fat splitting process. These components serve as potential secondary carbon sources or can be internalized to enhance the lipid content of the strain. β-oxidation-related genes were less active in SW_15_ (**Table S2**), but the higher expression of the 6 lipases (**Table S3**), especially *rtg2252* confirms a possible fatty acid degradation. This hints at the incorporation of exogenous fatty acids, which bypass β-oxidation and likely directly integrate into the cellular fatty acid pool through *de novo* lipid accumulation, contributing to an increase of lipid content. In fact, exogenous fatty acids contribute to upregulate Δ9 desaturase in *S. cerevisiae* (Bossie and Martin, 1989), consistent with the higher regulation of *rtg2309* (Δ9 desaturase) in SW_15_ **(Fig. 4)**. Furthermore, genes in “intracellular trafficking, secretion, and vesicular transport” are also upregulated, indicating active secretion/externalization activities in SW_15_. Fatty acids influx remain elusive, and it may occur via endocytosis, supported by the upregulation of *rtg1123* and *rtg2052* which encodes SLA1 and EPS15, proteins from the PAN1 complexes **(Fig. 3 and Table S1)**, which is involved in internalization of endosomes during actin-coupled endocytosis (Martin *et al*., 2007), possibly for incorporation of exogenous element and rapid modification of cell membrane as a response to stress (López-Hernández *et al*., 2020).

Moreover, we also found the possibility of sugar utilization in SW_15_, as glucosidases including *rtg4245* and *rtg428*, β-fructofuranosidase (*rtg769*) and glycolytic enzymes including putative rhamnogalacturonase (*rtg2468*) as well as a member of the glycosyl hydrolase family 88 (*rtg3298*) were upregulated **(Fig. 4)**. Among them, oligo-1,6-glucosidase (*rtg4245*) and β-fructofuranosidase (*rtg769*) were significantly upregulated, hence contributing the most to PCA1 **(Table S2)**. In addition, two MFS sugar transporters (*rtg1162* and *rtg5302*) were also upregulated **(Fig. 4 and Table S2)**. Simultaneously, *rtg2991* (FSP1) and *rtg3397* (STL1) responsible for glycerol uptake were downregulated, suggesting carbon catabolite repression (Bommareddy *et al*., 2017).

In terms of signaling, the most significant contrast between SW15 and Gly15 is observed in rtg3506, the activator of protein phosphatase 2A (PP2A), which exhibits a 16.5-fold higher expression level in SW15. This gene belongs to the phosphotyrosyl phosphatase activator (PTPA) family, known to promote PP2A activity. PP2A works in conjunction with the TORC pathway in yeast nitrogen sensing, but can also influence other cellular processes (Ariño *et al*., 2019). Hence, it is possible that TORC pathway is related to the upregulation of the genes involved in nitrogen recovery. The activation of β-oxidation by TORC is known as a response to nitrogen limitation (Gossing *et al*., 2018), and has also been reported in *R. toruloides* (Zhu *et al*., 2012).

Lastly, differences between SW_15_ and Gly_15_ may be influenced by pathways associated with cellular stress responses. Notably, there is an elevated expression of rtg4017 (RCK2), a gene known to be targeted by the high-osmolarity glycerol (HOG) pathway and involved in responding to both oxidative and osmotic stress (Bilsland-Marchesan *et al*., 2000). Additionally, the upregulation of *rtg4275* (RDS1) associated with stress response (Ludin *et al*., 1995), further suggests the involvement of stress-related pathways. In fact, both *rtg4017* and *rtg4275* has the highest expression in SW_15_, consistent with the associated lower biomass production. SW_15_ may also promote adaptive mechanisms, implying that the absence of a stress response in Gly_15_ could be attributed to the cells entering a dormant or quiescent state. Additionally, *rtg4659*, a heat shock transcription factor (HSF1), which is also a general stress effector, has similar expression levels in Gly_15_ and Gly_1.5_SC. While there is no significant difference in gene expression between SW_15_ and SW_1.5_SC, elevated expression of *rtg4659* in sweetwater as compared to Gly_15_/Gly_1.5_SC indicates that sweetwater can also trigger non-nutrient related stress responses even when it is diluted.

## 4. Conclusions

Utilization of crude sweetwater as a feedstock was evaluated for 21 lipid-accumulating yeasts. Improved growth was achieved by diluting and supplementing nutrients, but certain strains showed better lipid accumulation with crude sweetwater. Transcriptomics analysis of *R. toluroides* NRRL Y-6987, a top-performing strain, revealed a favorable C:N ratio in sweetwater for lipid accumulation. This study demonstrated the potential of sweetwater as a feedstock for microbial oil production, suggesting *R. toluroides* NRRL Y-6987 as a promising microbial oil producer. Insights obtained regarding key mechanisms of lipid accumulation induced by sweetwater also serves as a foundation for bioprocess optimization and strain engineering.

## CRediT authorship contribution statement

**Valériane Malika Keita:** Writing – original draft, Investigation, Formal analysis, Visualization

**Yi Qing Lee:** Writing – original draft, Formal analysis, Visualization

**Meiyappan Lakshmanan:** Formal analysis, Writing – Review & Editing

**Dave Siak-Wei Ow:** Investigation

**Paul Staniland:** Resources, Investigation

**Jessica Staniland:** Resources, Investigation

**Ian Savill:** Resources, Investigation

**Kang Lan Tee:** Conceptualization, Formal analysis

**Tuck Seng Wong:** Conceptualization, Formal analysis, Supervision, Writing – Review & Editing

**Dong-Yup Lee:** Conceptualization, Formal analysis, Supervision, Writing – Review & Editing Funding acquisition

## Acknowledgments

The research was supported by the Korea Innovation Foundation grant (2021-DD-UP-0369) funded by Ministry of Science and ICT and the Korea Institute of Planning and Evaluation for Technology in Food, Agriculture, Forestry and Fisheries (IPET) through High Value-added Food Technology Development Program (32136-05-1-HD050) funded by the MAFRA. VMK is supported by the Sheffield-A*STAR PhD scholarship. We also thank Dr. Pornkamol Unrean (BIOTEC, Thailand) for providing oleaginous yeast strains.

## Data availability

Data will be made available on request.

