## Appendix A. Supplementary data for "Sweetwater: an underrated crude glycerol for sustainable lipid production in non-conventional yeasts"

**Supplementary Figures**

**Fig. S1 Aspect and properties of the three types of sweetwater used in this study.** The asterisk indicates that SW_85_ was diluted before pH measurement.

**Fig. S2 Evaluating the glycerol tolerance of oleaginous yeasts** Cells were grown in 96-well microplates containing either refined glycerol diluted with synthetic complete nutrient mix for a final glycerol concentration of 2% (*w/v*) [black line: Gly_2_SC], 8% (*w/v*) [blue line: Gly_8_SC] or 16% (*w/v*) [red line: Gly_16_SC]. Cells were grown in quadruplicates and the averages OD_600_ and associated standard deviations were plotted against time.

**Fig. S3 Evaluating growth in different type of sweetwater diluted with the synthetic complete nutrient mic.** Cells were grown in 96-well microplates containing either in SW_15_ [black line: SW_15>1.5_SC], SW_35_ [teal line: SW_35>1.5_SC] or SW_85_ [purple line: SW_85>1.5_SC] diluted in synthetic complete nutrient mix for a final glycerol concentration of 1.5% (*w/v*). Cells were grown in quadruplicates and the averages OD_600_ and associated standard deviations were plotted against time.

**Fig. S4 Evaluating effect of sweetwater pH on growth.** Cells were grown in 96-well microplates containing either sweetwater with a 15% (*w/v*) glycerol content at native pH [black line: SW_15_ pH 3.95] or sweetwater with a 15% (*w/v*) glycerol content with a pH adjusted to 6 [teal line: SW_15_ pH 6.09]. Cells were grown in quadruplicates and the averages OD_600_ and associated standard deviations were plotted against time.

**Fig. S5 Clustergram of sample-to-sample Euclidean distances based on expression counts normalized by variance stabilizing transformation.**

**Fig. S6 Growth curves of *R. toruloides* NRRL Y-6987 in the 4 media used for the transcriptomic analysis.**

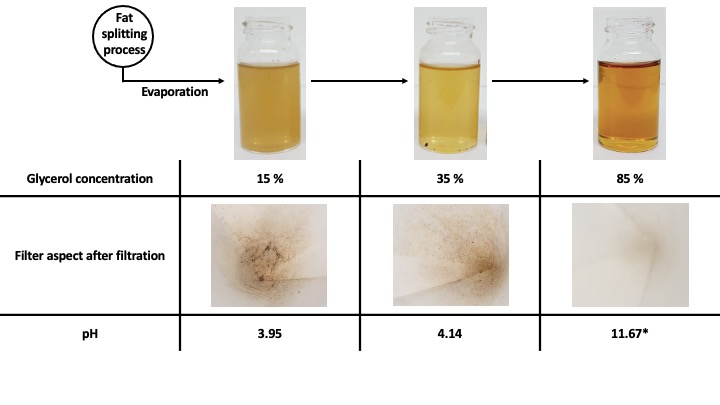

**Fig. S1**

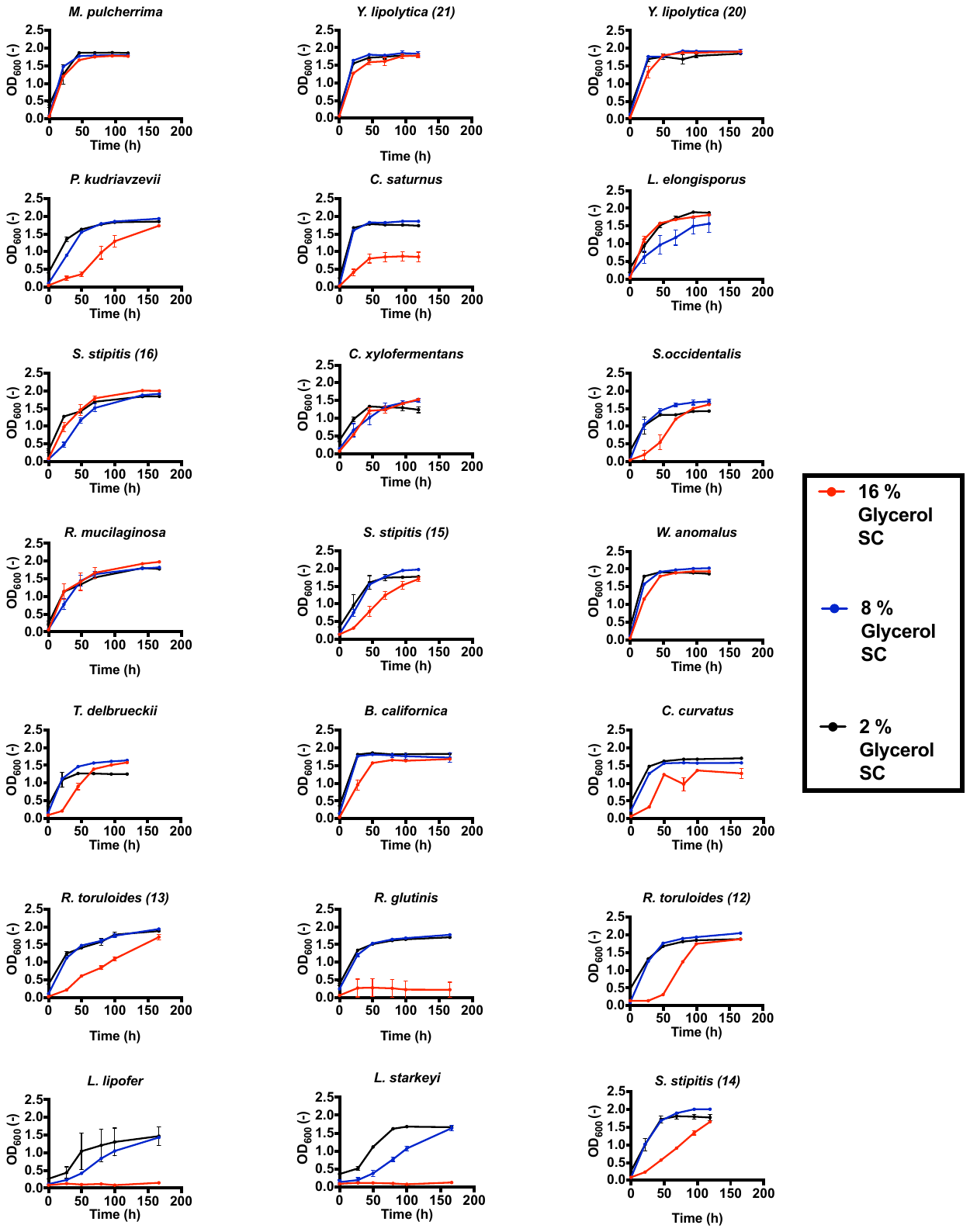

**Fig. S2**

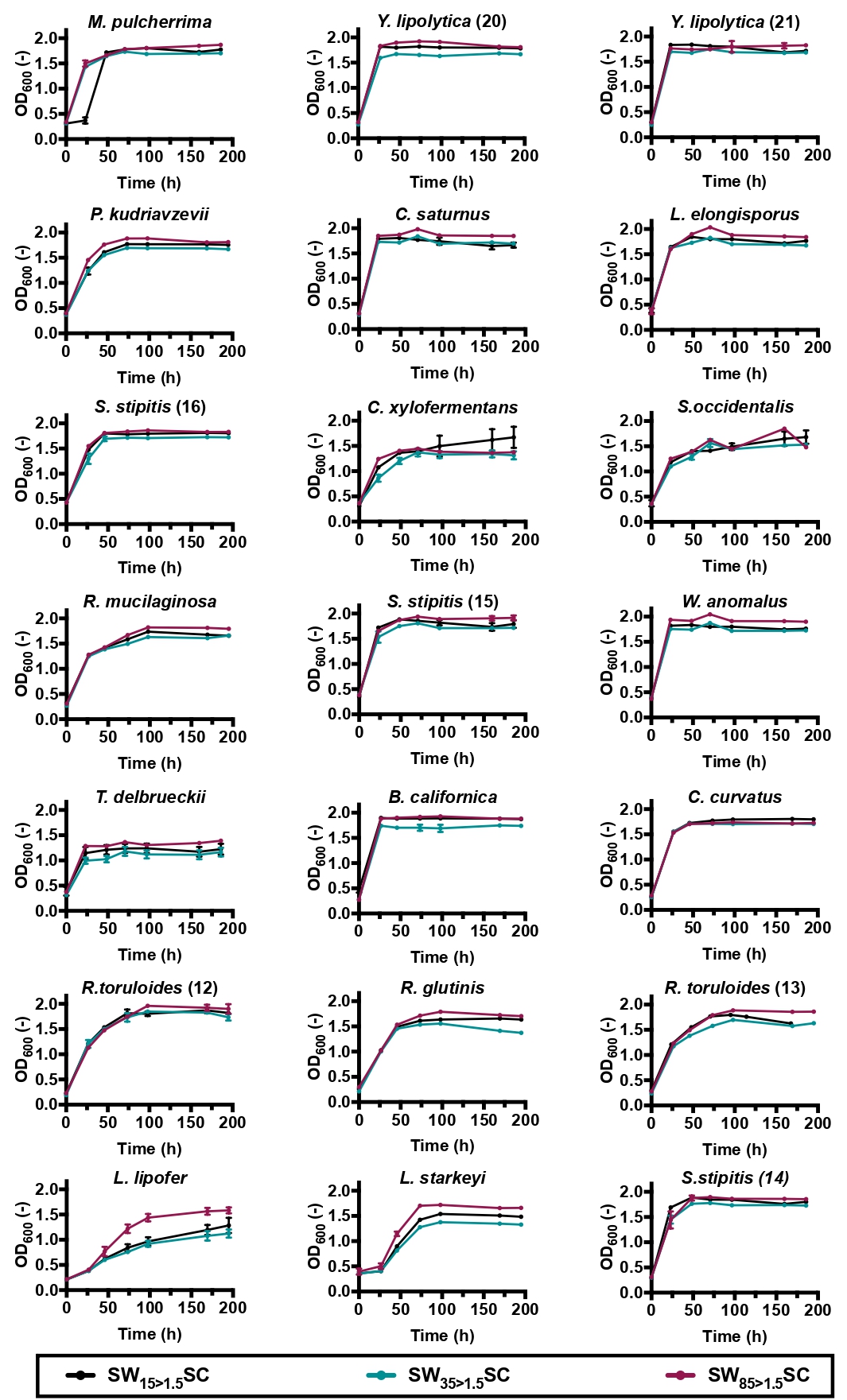

**Fig. S3**

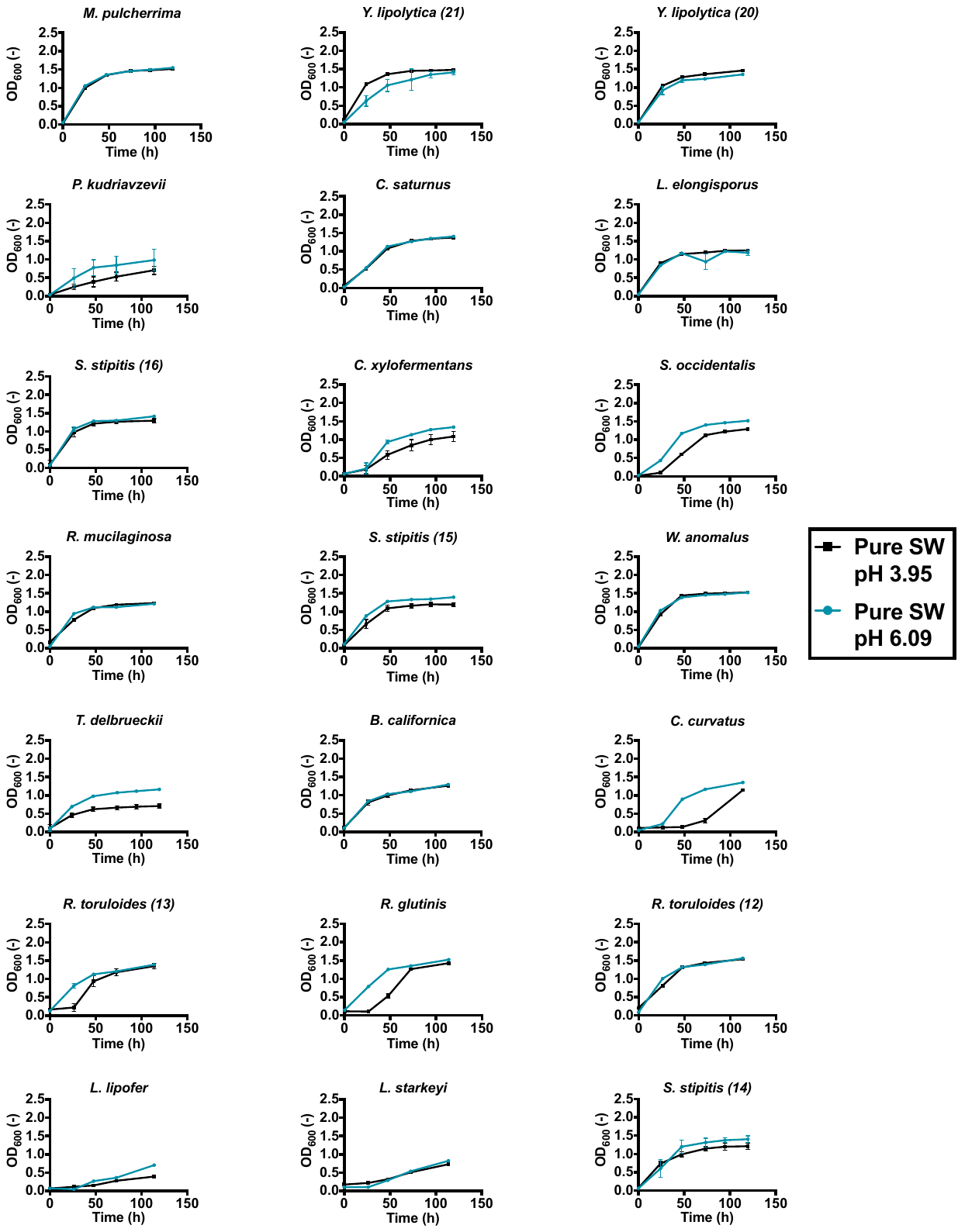

**Fig. S4**

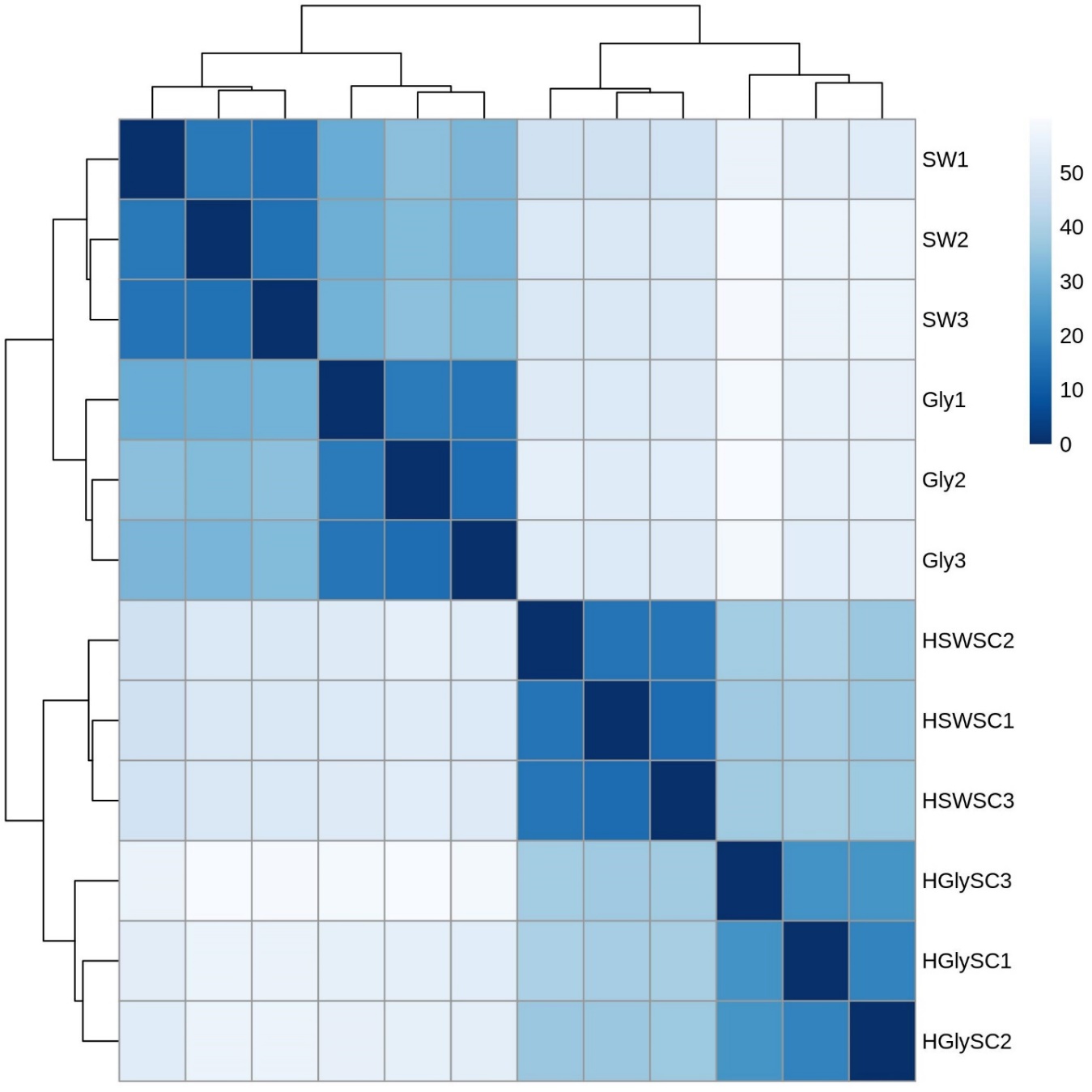

**Fig. S5**

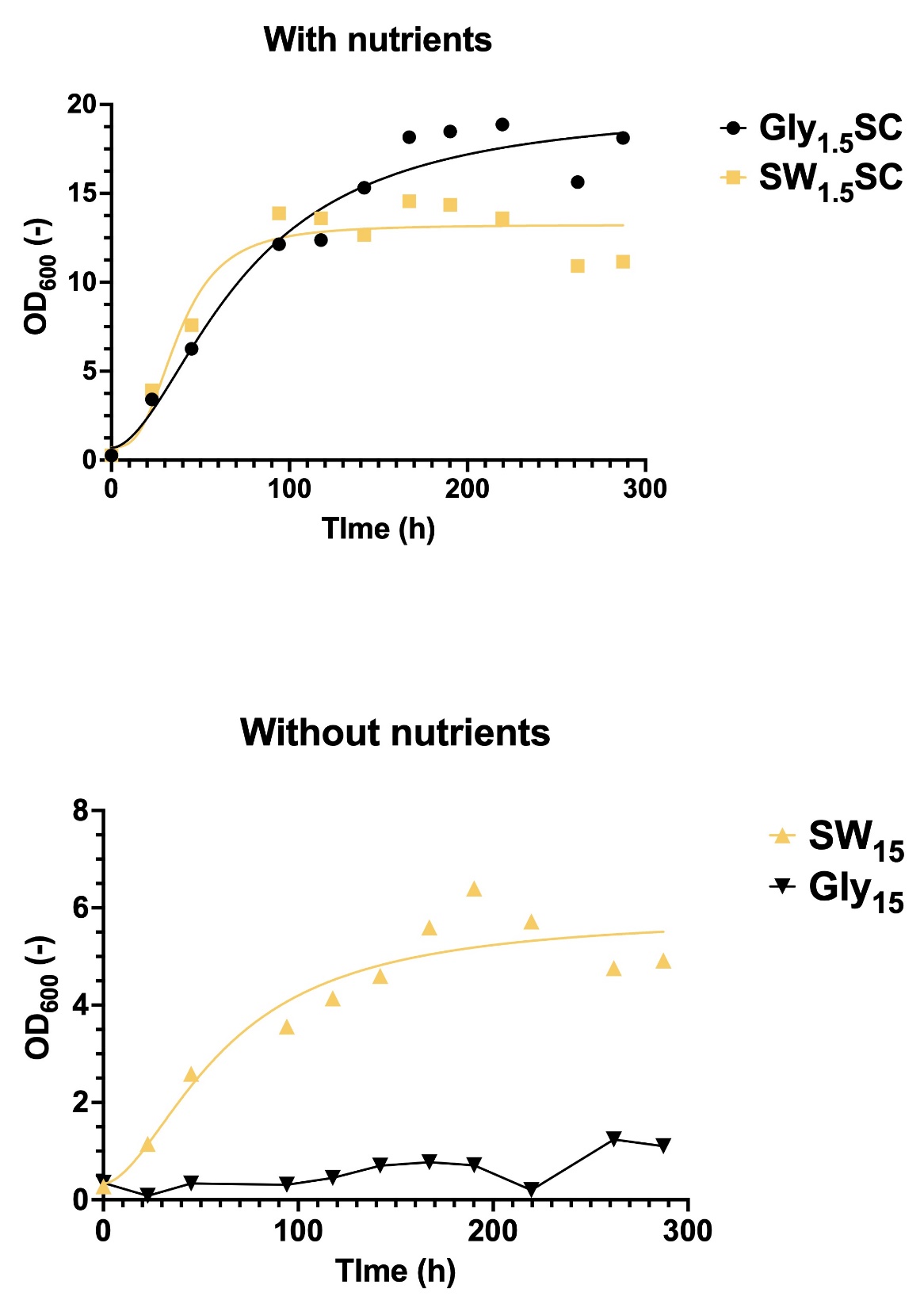

**Fig. S6**

**Supplementary Tables**

**Table S1 Annotation of genes represented in the figure 3*.***

**Table S2 Top 50 of annotated genes contributing the most to PC1 in the differential expression analysis of SW_15_ vs Gly_15_.** Gene ranks in terms of contribution to PC1 were given in the column Rank # and genes were sorted correspond sorted by descending contribution in each category. The smallest the rank is the highest the contribution will be. LFC (Log2 fold) gives the estimated change between gene expression level of in SW_15_ compared to Gly_15_ (Ctrl). This change is associated with a FDR adjusted p-value either non-significant (ns), non-computed (na), lower than 0.01 (<0.01), between 0.01 and 0.05 (<0.05) or between 0.05 and 0.1 (<0.1). Only padj <0.01 were considered significant in the main manuscript.

**Table S3 Differential expression results of SW_15_ vs Gly_15_ of relevant genes associated to lipid metabolism.** Gene ranks in terms of contribution to PC1 were given in the column Rank # and genes were sorted correspond sorted by descending contribution in each category. LFC (Log2 fold) gives the estimated change between gene expression level of in SW_15_ compared to Gly_15_ (Ctrl). This change is associated with a FDR adjusted p-value either non-significant (ns), non-computed (na), lower than 0.01 (<0.01), between 0.01 and 0.05 (<0.05) or between 0.05 and 0.1 (<0.1). Only padj <0.01 were considered significant in the main manuscript.

**Table S1**

| ID | KO | COG | Definition | Interclass |
| --- | --- | --- | --- | --- |
| rtg2117 | K01580 | E | glutamate decarboxylase [EC:4.1.1.15] | Amino acids transport and metabolism |
| rtg4590 | K08192 | G | MFS transporter, ACS family, allantoate permease | Carbohydrates transport and metabolism |
| rtg428 | K05349 | G | beta-glucosidase [EC:3.2.1.21] | Carbohydrates transport and metabolism |
| rtg1145 | - | G | Major Facilitator Superfamily | Carbohydrates transport and metabolism |
| rtg1762 | K08139 | G | MFS transporter, SP family, sugar:H+ symporter | Carbohydrates transport and metabolism |
| rtg3791 | K01183 | G | chitinase [EC:3.2.1.14] | Carbohydrates transport and metabolism |
| rtg4244 | K08141 | G | MFS transporter, SP family, general alpha glucoside:H+ symporter | Carbohydrates transport and metabolism |
| rtg4245 | K01182 | G | oligo-1,6-glucosidase [EC:3.2.1.10] | Carbohydrates transport and metabolism |
| rtg4615 | K02429 | G | MFS transporter, FHS family, L-fucose permease | Carbohydrates transport and metabolism |
| rtg4722 | K08178 | G | MFS transporter, SHS family, lactate transporter | Carbohydrates transport and metabolism |
| rtg5254 | K08139 | G | MFS transporter, SP family, sugar:H+ symporter | Carbohydrates transport and metabolism |
| rtg1110 | K02365 | D | separase [EC:3.4.22.49] | Cell cycle control, cell division, chromosome partitioning |
| rtg5072 | K11547 | D | kinetochore protein NDC80 | Cell cycle control, cell division, chromosome partitioning |
| rtg3275 | K03843 | M | alpha-1,3/alpha-1,6-mannosyltransferase [EC:2.4.1.132 2.4.1.257] | Cell wall/membrane/envelope biogenesis |
| rtg239 | K11253 | B | histone H3 | Chromatin structure and dynamics |
| rtg698 | K11251 | B | histone H2A | Chromatin structure and dynamics |
| rtg3504 | K11253 | - | histone H3 | Chromatin structure and dynamics\|Intracellular trafficking, secretion, and vesicular transport |
| rtg610 | K20520 | Z | drebrin-like protein | Cytoskeleton |
| rtg3437 | - | V | Male sterility protein | Defense mechanisms |
| rtg2031 | K00467 | C | lactate 2-monooxygenase [EC:1.13.12.4] | Energy production and conversion |
| rtg4853 | K00326;K17877 | C | cytochrome-b5 reductase [EC:1.6.2.2]\|nitrite reductase (NAD(P)H) [EC:1.7.1.4] | Energy production and conversion |
| rtg5031 | K00354 | C | NADPH2 dehydrogenase [EC:1.6.99.1] | Energy production and conversion |
| rtg1482 | K19792 | P | low-affinity ferrous iron transport protein | Inorganic ion transport and metabolism |
| rtg2366 | K03549;K17871 | P | KUP system potassium uptake protein\|NADH:ubiquinone reductase (non-electrogenic) [EC:1.6.5.9] | Inorganic ion transport and metabolism |
| rtg4153 | K20989 | P | urea-proton symporter | Inorganic ion transport and metabolism |
| rtg4852 | K02575 | P | MFS transporter, NNP family, nitrate/nitrite transporter | Inorganic ion transport and metabolism |
| rtg6000 | K03320 | P | ammonium transporter, Amt family | Inorganic ion transport and metabolism |
| rtg3429 | K18278 | - | Thiamine precursor biosynthesis enzyme | Metabolism of cofactors and vitamins |
| rtg6058 | K14709 | P | solute carrier family 39 (zinc transporter), member 1/2/3 | Inorganic ion transport and metabolism |
| rtg4154 | K20989 |  | urea-proton symporter | Intracellular trafficking, secretion, and vesicular transport |
| rtg3592 | K23516 | I | xanthine dioxygenase [EC:1.14.11.48] | Lipid transport and metabolism |
| rtg2138 | - | I | MaoC like domain (peroxisomal dehydratase - ncbiconfrimed) | Lipid transport and metabolism |
| rtg5959 | - | I | Acyl-CoA dehydrogenase, N-terminal domain | Lipid transport and metabolism |
| rtg4308 | K01487 | F | guanine deaminase [EC:3.5.4.3] | Nucleotide transport and metabolism |
| rtg4975 | K06910 |  | phosphatidylethanolamine-binding protein | Peptidases and inhibitors |
| rtg4874 | K03564 | O | peroxiredoxin Q/BCP [EC:1.11.1.15] | Post-translational modification and protein turnover |
| rtg5216 | K08770 | L | ubiquitin C | Replication, recombination and repair |
| rtg729 | - | A | RNA recognition motif. (a.k.a. RRM, RBD, or RNP domain) | RNA processing and modification |
| rtg206 | K13237 | Q | peroxisomal 2,4-dienoyl-CoA reductase [EC:1.3.1.34] | Secondary metabolites metabolism and transport |
| rtg2788 | K00002 | Q | alcohol dehydrogenase (NADP+) [EC:1.1.1.2] | Secondary metabolites metabolism and transport |
| rtg4053 | - | Q | Retinal pigment epithelial membrane protein | Secondary metabolites metabolism and transport |
| rtg4426 | K16216 | Q | benzil reductase ((S)-benzoin forming) [EC:1.1.1.320] | Secondary metabolites metabolism and transport |
| rtg6024 | - | Q | 2OG-Fe(II) oxygenase superfamily | Secondary metabolites metabolism and transport |
| rtg6040 | - | Q | short chain dehydrogenase reductase family protein | Secondary metabolites metabolism and transport |
| rtg6041 | - | Q | short chain dehydrogenase reductase family protein | Secondary metabolites metabolism and transport |
| rtg3996 | K00121 | Q | S-(hydroxymethyl)glutathione dehydrogenase / alcohol dehydrogenase [EC:1.1.1.284 1.1.1.1] | Secondary metabolites metabolism and transport |
| rtg3009 | K13953 | Q | alcohol dehydrogenase, propanol-preferring [EC:1.1.1.1] | Secondary metabolites metabolism and transport |
| rtg2375 | - | Q | Animal haem peroxidase | Secondary metabolites metabolism and transport |
| rtg814 | K06686;K08282 | T | cell cycle protein kinase DBF20 [EC:2.7.11.-]\|non-specific serine/threonine protein kinase [EC:2.7.11.1] | Signal transduction mechanisms |
| rtg5735 | K11593 | J | eukaryotic translation initiation factor 2C | Translation, ribosomal structure and biogenesis |

**Table S2**

| ID | Rank # | LogFC | p_adj_ | KO | COG | Definition |
| --- | --- | --- | --- | --- | --- | --- |
| Nitrogen scavenging/Nitrogen recycling | | | | | | |
| rtg4852 | 10 | -2.19279 | <0.01 | K02575 | P | MFS transporter, NNP family, nitrate/nitrite transporter |
| rtg3792 | 33 | 2.216161 | <0.01 | K01426 | E | Amidase [EC:3.5.1.4] |
| rtg3315 | 38 | 1.47717 | <0.01 |  | T | OPT oligopeptide transporter protein |
| rtg4853 | 44 | -1.43723 | <0.01 | K00326;K17877 | C | Cytochrome-b5 reductase [EC:1.6.2.2]\|nitrite reductase (NAD(P)H) [EC:1.7.1.4] |
| rtg5673 | 45 | 1.560374 | <0.01 | K03305 | E | Proton-dependent oligopeptide transporter, POT family |
| Signaling | | | | | | |
| rtg3506 | 7 | 4.04169 | <0.01 | K17605 | D;T | Serine/threonine-protein phosphatase 2A activator |
| rtg4017 | 24 | 1.723819 | <0.01 | K23785 | T | Serine/threonine-protein kinase RCK2 [EC:2.7.11.1] |
| rtg4275 | 39 | 1.50702 | <0.01 |  | S | Stress response protein Rds1 |
| rtg4659 | 59 | 1.451221 | <0.01 | K09419 | K | Heat shock transcription factor, other eukaryote |
| rtg4708 | 26 | 1.621092 | <0.01 |  | T | Cytoplasm protein (RGC2) |
| Membrane remodeling | | | | | | |
| rtg3898 | 8 | -5.3022 | <0.01 |  |  | Rare lipoprotein A (rlpa)-like double-psi beta-barrel/expansin |
| rtg6092 | 9 | 3.854252 | <0.01 |  | O | Fasciclin domain (involved in cell adhesion)/Beta-ig-h3 fasciclin |
| rtg2052 | 43 | 1.484737 | <0.01 | K12472 | T;U | Epidermal growth factor receptor substrate 15 |
| rtg1123 | 60 | 1.359099 | <0.01 | K20046 | Z | Actin cytoskeleton-regulatory complex protein SLA1 |
| Protein modification and turnover | | | | | | |
| rtg5563 | 15 | 2.083641 | <0.01 | K01875 | J | Seryl-trna synthetase [EC:6.1.1.11] |
| rtg3923 | 42 | 1.524287 | <0.01 | K03695 | O | ATP-dependent Clp protease ATP-binding subunit clpb |
| rtg2372 | 63 | 1.347649 | <0.01 |  | O | ATP-dependent protease La (LON) domain |
| Energy and redox balance | | | | | | |
| rtg5547 | 6 | 3.209785 | <0.01 | K00228 | H | Coproporphyrinogen III oxidase [EC:1.3.3.3] |
| rtg5031 | 32 | 1.59539 | <0.01 | K00354 | C | NADPH2 dehydrogenase [EC:1.6.99.1] |
| rtg2788 | 37 | 1.509143 | <0.01 | K00002 | Q | Alcohol dehydrogenase (NADP+) [EC:1.1.1.2] |
| Carbon catabolite repression | | | | | | |
| rtg4245 | 1 | 3.477264 | <0.01 | K01182 | G | Oligo-1,6-glucosidase [EC:3.2.1.10] |
| rtg5013 | 13 | 1.949483 | <0.01 | K01621 | G | Xylulose-5-phosphate/fructose-6-phosphate phosphoketolase [EC:4.1.2.9 4.1.2.22] |
| rtg769 | 14 | 1.93709 | <0.01 | K01193 | G | Beta-fructofuranosidase [EC:3.2.1.26] |
| rtg5302 | 19 | 1.801282 | <0.01 | K08139 | G | MFS transporter, SP family, sugar:H+ symporter (HXT family) |
| rtg428 | 25 | 2.701482 | <0.01 | K05349 | G | Beta-glucosidase [EC:3.2.1.21] |
| rtg2468 | 27 | 1.702245 | <0.01 |  |  | Pbh1 domain (= tertiary structures of pectate lyases and rhamnogalacturonase A) |
| rtg1823 | 29 | 1.788605 | <0.01 |  | E;G | Triose-phosphate Transporter family |
| rtg1162 | 34 | 1.699774 | <0.01 |  | G | Major facilitator superfamily |
| rtg3298 | 56 | 1.396897 | <0.01 |  | S | Glycosyl hydrolase family 88 |
| Fatty acid uptake | | | | | | |
| rtg2252 | 31 | 2.678024 | <0.01 |  |  | Lipase B precursor |
| rtg2309 | 46 | 1.418252 | <0.01 | K00507 | I | Stearoyl-coa desaturase (Delta-9 desaturase) [EC:1.14.19.1] |
| Beta-oxidation and Glyoxylate cycle related | | | | | | |
| rtg2130 | 3 | -5.98348 | n.a. | K01638 | C | Malate synthase [EC:2.3.3.9] |
| rtg5452 | 41 | -1.53047 | <0.01 |  | I | Acyl-coa dehydrogenase, C-terminal domain |
| rtg4139 | 64 | -1.74603 | <0.01 | K00624 | I | Carnitine O-acetyltransferase [EC:2.3.1.7] |
| Fatty synthesis | | | | | | |
| rtg2312 | 17 | 1.797549 | <0.01 | K01648 | C | ATP citrate (pro-S)-lyase [EC:2.3.3.8] |
| rtg127 | 50 | 1.355341 | <0.01 | K00667 | I | Fatty acid synthase subunit alpha, fungi type [EC:2.3.1.86] |
| rtg226 | 57 | 1.329832 | <0.01 | K11262 | I | Acetyl-coa carboxylase / biotin carboxylase 1 [EC:6.4.1.2 6.3.4.14 2.1.3.15] |
| rtg204 | 58 | 1.376327 | <0.01 | K00667;K00668 | I | Fatty acid synthase subunit beta, fungi type [EC:2.3.1.86] |
| Unclassified | | | | | | |
| rtg5628 | 2 | 2.609292 | <0.01 |  |  | Stigma-specific protein, Stig1 |
| rtg3278 | 4 | 3.410903 | <0.01 |  | Q | ABC-2 type transporter |
| rtg4452 | 5 | 4.020687 | <0.01 |  | S | Inherit from basnog: delayed-type hypersensitivity antigen |
| rtg3026 | 21 | -2.24292 | <0.01 |  | S | Protein similar to cwfj C-terminus 2 |
| rtg2410 | 28 | 2.059587 | <0.01 |  | G | Fungal trichothecene (sesquiterpenoids) efflux pump (MFS familly) |
| rtg2948 | 30 | 2.616607 | <0.01 |  | S | Inherit from funog: CCCH zinc finger DNA binding protein |
| rtg2495 | 35 | 1.401479 | ns | | S | Predicted membrane protein (DUF2231) |
| rtg617 | 40 | -2.3205 | <0.01 | K00622 | Q | Arylamine N-acetyltransferase [EC:2.3.1.5] |
| rtg5922 | 48 | -2.1786 | <0.01 |  | S | SCP (Cystein-rich secretory protein) |
| rtg3821 | 53 | 1.752634 | <0.01 |  | S | FAD dependent oxidoreductase |
| rtg3945 | 54 | 1.42675 | <0.01 |  | S | Zinc-binding dehydrogenase |
| rtg5476 | 61 | 1.64211 | <0.01 |  | Q | ABC transporter transmembrane region |

**Table S3**

| ID | Rank # | LogFC | p_adj_ | KO | COG | Definition |
| --- | --- | --- | --- | --- | --- | --- |
| Acetyl-CoA metabolism | | | | | | |
| Acetyl-CoA synthetase | | | | | | |
| rtg931 | 1711 | 0.463515 | <0.05 | K01895 | H | Acetyl-coa synthetase [EC:6.2.1.1] |
| ATP:Citrate lyase | | | | | | |
| rtg2312 | 17 | 1.797549 | <0.01 | K01648 | C | ATP citrate (pro-S)-lyase [EC:2.3.3.8] |
| Acetyl-CoA C-acetyltransferase (ACAT) | | | | | | |
| rtg1529 | 3125 | -0.22239 | ns | K07513 | I | Acetyl-coa acyltransferase 1 [EC:2.3.1.16] |
| rtg3157 | 4703 | -0.11376 | ns | K07513 | I | Acetyl-coa acyltransferase 1 [EC:2.3.1.16] |
| rtg4298 | 1766 | -0.36792 | <0.01 | K07513 | I | Acetyl-coa acyltransferase 1 [EC:2.3.1.16] |
| Citrate synthase | | | | | | |
| rtg1700 | 1590 | 0.529734 | <0.05 | K01647 | C | Citrate synthase [EC:2.3.3.1] |
| rtg2745 | 3093 | 0.528872 | ns | K01647 | C | Citrate synthase [EC:2.3.3.1] |
| rtg2747 | 251 | 0.908961 | <0.01 | K01647 | C | Citrate synthase [EC:2.3.3.1] |
| Malate dehydrogenases | | | | | | |
| rtg1361 | 1394 | 0.429893 | <0.01 | K00026 | C | Malate dehydrogenase [EC:1.1.1.37] |
| rtg1596 | 2036 | 0.43835 | ns | K00027 | C | Malate dehydrogenase (oxaloacetate-decarboxylating) [EC:1.1.1.38] |
| rtg3452 | 359 | 0.777535 | <0.01 | K00026 | C | Malate dehydrogenase [EC:1.1.1.37] |
| rtg6104 | 649 | -1.02196 | <0.01 | K00027;K00029 | C | Malate dehydrogenase (oxaloacetate-decarboxylating) [EC:1.1.1.38]\|malate dehydrogenase (oxaloacetate-decarboxylating)(NADP+) [EC:1.1.1.40] |
| Fumarase | | | | | | |
| rtg5411 | 1021 | 0.559453 | <0.01 | K01679 | C | Fumarate hydratase, class II [EC:4.2.1.2] |
| Succinate dH | | | | | | |
| rtg1051 | 663 | 0.683249 | <0.01 | K18168 | S | Succinate dehydrogenase assembly factor 2 |
| rtg1256 | 286 | 0.828923 | <0.01 | K00234 | C | Succinate dehydrogenase (ubiquinone) flavoprotein subunit [EC:1.3.5.1] |
| rtg1876 | 2552 | -0.45795 | ns | K18167 | O | Succinate dehydrogenase assembly factor 1 |
| rtg4479 | 1105 | 0.505001 | <0.01 | K00236 | C | Succinate dehydrogenase (ubiquinone) cytochrome b560 subunit |
| rtg5261 | 1913 | 0.340104 | <0.05 | K00235 | C | Succinate dehydrogenase (ubiquinone) iron-sulfur subunit [EC:1.3.5.1] |
| rtg603 | 987 | 0.600242 | <0.01 | K00237 | C | Succinate dehydrogenase (ubiquinone) membrane anchor subunit |
| Succinyl-CoA synthase | | | | | | |
| rtg5615 | 1600 | 0.385984 | <0.01 | K01900 | C | Succinyl-coa synthetase beta subunit [EC:6.2.1.4 6.2.1.5] |
| rtg5749 | 2564 | 0.255147 | <0.05 | K01899 | C | Succinyl-coa synthetase alpha subunit [EC:6.2.1.4 6.2.1.5] |
| alpha-ketoglutarate dH | | | | | | |
| rtg1066 | 324 | 0.83069 | <0.01 | K00658 | C | 2-oxoglutarate dehydrogenase E2 component (dihydrolipoamide succinyltransferase) [EC:2.3.1.61] |
| rtg2808 | 180 | 0.972097 | <0.01 | K00164 | G | 2-oxoglutarate dehydrogenase E1 component [EC:1.2.4.2] |
| rtg3479 | 4498 | 0.145618 | ns | K00164;K15791 | G | 2-oxoglutarate dehydrogenase E1 component [EC:1.2.4.2]\|probable 2-oxoglutarate dehydrogenase E1 component DHKTD1 [EC:1.2.4.2] |
| Isocitrate dH | | | | | | |
| rtg3415 | 3893 | 0.145291 | ns | K00031 | C | Isocitrate dehydrogenase [EC:1.1.1.42] |
| rtg5809 | 1036 | 0.495925 | <0.01 | K00030 | E | Isocitrate dehydrogenase (NAD+) [EC:1.1.1.41] |
| rtg5810 | 418 | 0.772541 | <0.01 | K00030 | E | Isocitrate dehydrogenase (NAD+) [EC:1.1.1.41] |
| Aconitase | | | | | | |
| rtg4484 | 252 | 0.846646 | <0.01 | K01681 | C | Aconitate hydratase [EC:4.2.1.3] |
| Isocitrate lyase | | | | | | |
| rtg5517 | 3932 | 0.126558 | ns | K01637 | C | Isocitrate lyase [EC:4.1.3.1] |
| Malate synthase | | | | | | |
| rtg2130 | 3 | -5.98348 | n.a | K01638 | C | Malate synthase [EC:2.3.3.9] |
| rtg3633 | 136 | 1.127702 | <0.01 | K01638 | C | Malate synthase [EC:2.3.3.9] |
| De novo lipid synthesis | | | | | | |
| Acetyl-CoA carboxylase (ACC) | | | | | | |
| rtg226 | 57 | 1.329832 | <0.01 | K11262 | I | Acetyl-coa carboxylase / biotin carboxylase 1 [EC:6.4.1.2 6.3.4.14 2.1.3.15] |
| Fatty acid synthase (FAS) | | | | | | |
| rtg127 | 50 | 1.355341 | <0.01 | K00667 | I | Fatty acid synthase subunit alpha, fungi type [EC:2.3.1.86] |
| rtg204 | 58 | 1.376327 | <0.01 | K00667;K00668 | I | Fatty acid synthase subunit beta, fungi type [EC:2.3.1.86] |
| 1-acylglycerol-3-phosphate O-acyltransferase (AGPAT) | | | | | | |
| rtg5664 | 2596 | 0.266996 | <0.1 | K13519 | I | Lysophospholipid acyltransferase [EC:2.3.1.51 2.3.1.23 2.3.1.-] |
| rtg2173 | 5298 | -0.02788 | ns | | I | Acyltransferase (lysophospholipid acyltransferase) |
| rtg5355 | 5810 | -0.00527 | ns | | I | Acyltransferase (lysophosphatidic acid acyltransferase / lysophosphatidylinositol acyltransferase) |
| Glycerol-3-phosphate acyltransferase (GPAT) | | | | | | |
| rtg1283 | 3535 | -0.25377 | ns | K13507 | I | Glycerol-3-phosphate O-acyltransferase / dihydroxyacetone phosphate acyltransferase [EC:2.3.1.15 2.3.1.42] |
| rtg3012 | 801 | 0.567038 | <0.01 | K13507 | I | Glycerol-3-phosphate O-acyltransferase / dihydroxyacetone phosphate acyltransferase [EC:2.3.1.15 2.3.1.42] |
| rtg3414 | 4623 | -0.1619 | ns | K13507 | I | Glycerol-3-phosphate O-acyltransferase / dihydroxyacetone phosphate acyltransferase [EC:2.3.1.15 2.3.1.42] |
| Phosphatidic acid phosphatase (PPAP) | | | | | | |
| rtg1218 | 3731 | 0.174715 | ns | K18693 | I | Diacylglycerol diphosphate phosphatase / phosphatidate phosphatase [EC:3.1.3.81 3.1.3.4] |
| rtg3174 | 2980 | 0.923743 | ns | |  | (refseq) bifunctional diacylglycerol diphosphate phosphatase/phosphatidate phosphatase<\|acidppc |
| rtg3837 | 4851 | 0.398036 | ns | |  |  |
| Diacylglycerol O-acyltransferase (DGAT) and associated | | | | | | |
| rtg257 | 4851 | 0.530466 | <0.1 | K15728 | I | Phosphatidate phosphatase LPIN [EC:3.1.3.4] |
| rtg267 | 1385 | 0.478952 | <0.05 | K00679 | I | Phospholipid:diacylglycerol acyltransferase [EC:2.3.1.158] |
| alpha, omega-oxidation | | | | | | |
| Aldehyde dehydrogenase | | | | | | |
| rtg1234 | 3133 | 0.237564 | ns | K00128;K00129 | C | Aldehyde dehydrogenase (NAD(P)+) [EC:1.2.1.5]\|aldehyde dehydrogenase (NAD+) [EC:1.2.1.3] |
| rtg1331 | 1672 | -0.61742 | <0.05 |  | C | Aldehyde dehydrogenase family |
| rtg1455 | 1741 | 0.41167 | <0.05 |  | V | D-lactaldehyde dehydrogenase |
| rtg3413 | 4758 | 0.105004 | ns | K00128 | C | Aldehyde dehydrogenase (NAD+) [EC:1.2.1.3] |
| rtg4159 | 1901 | 0.360954 | <0.01 | K00128;K14085 | C | Aldehyde dehydrogenase (NAD+) [EC:1.2.1.3]\|aldehyde dehydrogenase family 7 member A1 [EC:1.2.1.31 1.2.1.8 1.2.1.3] |
| rtg4562 | 976 | -0.64903 | <0.01 | K00128 | C | Aldehyde dehydrogenase (NAD+) [EC:1.2.1.3] |
| rtg4984 | 668 | -0.64935 | <0.01 | K00128 | C | Aldehyde dehydrogenase (NAD+) [EC:1.2.1.3] |
| rtg513 | 4741 | 0.101235 | ns | K00128 | C | Aldehyde dehydrogenase (NAD+) [EC:1.2.1.3] |
| rtg5244 | 2097 | -0.84413 | <0.05 | K00128 | C | Aldehyde dehydrogenase (NAD+) [EC:1.2.1.3] |
| rtg738 | 1018 | -0.50214 | <0.01 | K00128 | C | Aldehyde dehydrogenase (NAD+) [EC:1.2.1.3] |
| Alcohol dehydrogenase | | | | | | |
| rtg1625 | 1196 | 0.602372 | <0.01 | K00002 | Q | Alcohol dehydrogenase (NADP+) [EC:1.1.1.2] |
| rtg2078 | 144 | 1.105919 | <0.01 | K00002 | Q | Alcohol dehydrogenase (NADP+) [EC:1.1.1.2] |
| rtg2788 | 37 | 1.509143 | <0.01 | K00002 | Q | Alcohol dehydrogenase (NADP+) [EC:1.1.1.2] |
| rtg289 |  | 0.795027 | <0.01 | K13953 | Q | Alcohol dehydrogenase, propanol-preferring [EC:1.1.1.1] |
| rtg2985 | 4952 | -0.17527 | ns | K13953 | Q | Alcohol dehydrogenase, propanol-preferring [EC:1.1.1.1] |
| rtg2999 | 114 | 1.40059 | <0.01 |  | C | Aryl-alcohol dehydrogenase |
| rtg3009 | 844 | 1.106993 | <0.01 | K13953 | Q | Alcohol dehydrogenase, propanol-preferring [EC:1.1.1.1] |
| rtg3974 | 2441 | -0.45701 | <0.05 | K13954 | C | Alcohol dehydrogenase [EC:1.1.1.1] |
| rtg3996 | 776 | 0.554074 | <0.01 | K00121 | Q | S-(hydroxymethyl)glutathione dehydrogenase / alcohol dehydrogenase [EC:1.1.1.284 1.1.1.1] |
| rtg4272 | 4812 | -0.14568 | ns | K13954 | C | Alcohol dehydrogenase [EC:1.1.1.1] |
| rtg4461 | 5004 | -0.12369 | ns | K13953 | Q | Alcohol dehydrogenase, propanol-preferring [EC:1.1.1.1] |
| rtg4464 | 4973 | 0.163424 | ns | K13953 | Q | Alcohol dehydrogenase, propanol-preferring [EC:1.1.1.1] |
| rtg5549 | 2598 | 0.34862 | ns | | C | Alcohol dehydrogenase groes-like domain |
| rtg6061 | 5183 | -0.07196 | ns | K13953;K13955 | C | Alcohol dehydrogenase, propanol-preferring [EC:1.1.1.1]\|zinc-binding alcohol dehydrogenase/oxidoreductase |
| rtg6101 | 5370 | -0.2791 | ns | | Q | Alcohol dehydrogenase groes-like domain |
| Cytochrome P450 alkane monooxygenase/cytochrome P450 reductase complex (CYP + CPR) | | | | | | |
| rtg3373 | 2175 | -0.77424 | <0.05 |  | Q | Cytochrome P450 |
| rtg3536 | 1733 | -0.5131 | <0.1 |  | Q | Cytochrome p450 |
| rtg4089 | 380 | 0.369566 | ns | | Q | Cytochrome p450 |
| rtg5028 | 2606 | 0.276924 | ns | | Q | Cytochrome P450 |
| Lipases | | | | | | |
| rtg2110 | 3745 | 0.321266 | ns | K17756 | O | Secretory lipase (family LIP) |
| rtg2248 | 4610 | -0.09527 | ns | K06130;K06999 | | Lysophospholipase II [EC:3.1.1.5]\|phospholipase/carboxylesterase |
| rtg2252 | 31 | 2.678024 | <0.01 |  |  | (refseq) putative Lipase B precursor< |
| rtg2257 | 4026 | 0.806034 | ns | | I | Inherit from funog: GDSL-like Lipase/Acylhydrolase |
| rtg2311 | 399 | 1.074245 | <0.01 | K01054 | I | Acylglycerol lipase [EC:3.1.1.23] |
| rtg2505 | 1516 | 0.555816 | <0.05 | K01046 | I | Triacylglycerol lipase [EC:3.1.1.3] |
| rtg2599 | 1517 | 0.555816 | <0.05 | K01046 | I | Triacylglycerol lipase [EC:3.1.1.3] |
| rtg3022 | 2988 | 0.242132 | ns | K14018 | I | Phospholipase A-2-activating protein |
| rtg3516 | 4122 | 0.146469 | ns | | I;O;T | Lipase (class 3) |
| rtg3673 | 538 | 1.268124 | <0.01 |  | O | Secretory lipase |
| rtg3917 | 101 | 1.41606 | <0.01 | K13333 | I | Lysophospholipase [EC:3.1.1.5] |
| rtg409 | 3139 | 0.459692 | ns | | T | Extracellular triacylglycerol lipase precursor (EC 3.1.1.3) |
| rtg434 | 2261 | -0.49125 | <0.1 | K13985 | S | N-acyl-phosphatidylethanolamine-hydrolysing phospholipase D [EC:3.1.4.54] |
| rtg4451 | 84 | 1.900333 | <0.01 | K13278 | E | 60kda lysophospholipase [EC:3.1.1.5 3.1.1.47 3.5.1.1] |
| rtg4582 | 2328 | 0.254256 | ns | K01046;K15979 | K | Staphylococcal nuclease domain-containing protein 1\|triacylglycerol lipase [EC:3.1.1.3] |
| rtg4806 | 1948 | 0.372918 | <0.1 |  |  | (refseq) triglyceride lipase< |
| rtg4919 | 4041 | 0.187818 | ns | | I | GDSL-like Lipase/Acylhydrolase |
| rtg498 | 2099 | -0.50138 | <0.1 | K01115;K15429 | | Phospholipase D1/2 [EC:3.1.4.4]\|trna (guanine37-N1)-methyltransferase [EC:2.1.1.228] |
| rtg511 | 5730 | 0.109941 | ns | K06130;K06999 | I | Lysophospholipase II [EC:3.1.1.5]\|phospholipase/carboxylesterase |
| rtg5423 | 2591 | 0.237573 | ns | K01115 | I | Phospholipase D1/2 [EC:3.1.4.4] |
| rtg5514 | 2729 | -0.37738 | ns | K01054 | I | Acylglycerol lipase [EC:3.1.1.23] |
| rtg5578 | 831 | 0.670736 | <0.01 | K01052 | I | Lysosomal acid lipase/cholesteryl ester hydrolase [EC:3.1.1.13] |
| rtg6015 | 5646 | -0.0113 | ns | | I | Phospholipase/carboxylesterase |
| rtg6067 | 5302 | 0.022295 | ns | K00997;K13333;K16342 | I | Cytosolic phospholipase A2 [EC:3.1.1.4]\|holo-[acyl-carrier protein] synthase [EC:2.7.8.7]\|lysophospholipase [EC:3.1.1.5] |
| rtg913 | 3449 | -0.18737 | ns | | I | Partial alpha/beta-hydrolase lipase region |
| β-oxidation | | | | | | |
| Acyl-Coa dehydrogenase | | | | | | |
| rtg2045 | 1273 | -1.29835 | <0.01 |  |  | (refseq) bifunctional hydroxyacyl-coa dehydrogenase/enoyl-coa hydratase FOX2<\|short chain dehydrogenase |
| rtg23 | 2267 | -0.86759 | <0.05 |  | I | Acyl-coa dehydrogenase |
| rtg2805 | 4708 | -0.12365 | ns | K00253 | I | Isovaleryl-coa dehydrogenase [EC:1.3.8.4] |
| rtg4074 | 5070 | -0.08105 | ns | | I | Acyl-coa dehydrogenase, C-terminal domain |
| rtg4571 | 2264 | 0.513542 | <0.05 | K00249 | I | Acyl-coa dehydrogenase [EC:1.3.8.7] |
| rtg5452 | 41 | -1.53047 | <0.01 |  | I | Acyl-coa dehydrogenase, C-terminal domain |
| rtg5959 | 2923 | -0.23662 | ns | | I | Acyl-coa dehydrogenase, N-terminal domain |
| rtg701 | 669 | -0.70349 | <0.01 |  | I | Acyl-coa dehydrogenase, C-terminal domain |
| enoyl-CoA hydratase | | | | | | |
| rtg2045 | 1273 | -1.29835 | <0.01 |  |  | (refseq) bifunctional hydroxyacyl-coa dehydrogenase/enoyl-coa hydratase FOX2<\|short chain dehydrogenase |
| rtg3404 | 863 | -0.60749 | <0.01 |  | I | Enoyl-coa hydratase/isomerase family |
| rtg5742 | 2945 | -0.73283 | ns | K05605;K05607 | I | 3-hydroxyisobutyryl-coa hydrolase [EC:3.1.2.4]\|methylglutaconyl-coa hydratase [EC:4.2.1.18] |
| rtg781 | 4023 | -0.15761 | ns | K05605;K07511 | I | 3-hydroxyisobutyryl-coa hydrolase [EC:3.1.2.4]\|enoyl-coa hydratase [EC:4.2.1.17] |
| 3L-hydroxyacyl-coa dehydrogenases | | | | | | |
| rtg3194 | 848 | -1.04956 | <0.01 | K00074 | I | 3-hydroxybutyryl-coa dehydrogenase [EC:1.1.1.157] |
| rtg1254 | 5159 | 0.077437 | ns | K08683 |  | 3-hydroxyacyl-coa dehydrogenase / 3-hydroxy-2-methylbutyryl-coa dehydrogenase [EC:1.1.1.35 1.1.1.178] |
| β-ketothiolase | | | | | | |
| rtg1529 | 3125 | -0.22239 | ns | K07513 | I | Acetyl-coa acyltransferase 1 [EC:2.3.1.16] |
| rtg3157 | 4703 | -0.11376 | ns | K07513 | I | Acetyl-coa acyltransferase 1 [EC:2.3.1.16] |
| rtg4298 | 1766 | -0.36792 | <0.01 | K07513 | I | Acetyl-coa acyltransferase 1 [EC:2.3.1.16] |
| Caroteinoids synthesis | | | | | | |
| rtg281 | 1189 | 0.55891 | <0.01 | K00804 | H | Geranylgeranyl diphosphate synthase, type III [EC:2.5.1.1 2.5.1.10 2.5.1.29] |
| rtg4052 | 1768 | 0.356349 | <0.1 | K17841 | I | 15-cis-phytoene synthase / lycopene beta-cyclase [EC:2.5.1.32 5.5.1.19] |
| rtg4054 | 337 | 0.801796 | <0.01 | K15745 | H | Phytoene desaturase (3,4-didehydrolycopene-forming) [EC:1.3.99.30] |
| rtg1071 | 2788 | 0.252253 | ns | K01641 | I | Hydroxymethylglutaryl-coa synthase [EC:2.3.3.10] |
| rtg2210 | 1378 | 0.499395 | <0.1 | K00021 | I | Hydroxymethylglutaryl-coa reductase (NADPH) [EC:1.1.1.34] |
| rtg64 | 2091 | -0.42366 | <0.05 | K01823 | P | Isopentenyl-diphosphate Delta-isomerase [EC:5.3.3.2] |
| rtg55 | 6078 | 0.072335 | ns | K01823 | Q | Isopentenyl-diphosphate Delta-isomerase [EC:5.3.3.2] |
| rtg281 | 1189 | 0.55891 | <0.01 | K00804 | H | Geranylgeranyl diphosphate synthase, type III [EC:2.5.1.1 2.5.1.10 2.5.1.29] |
| rtg5984 | 2166 | -0.35186 | ns | K00787 | H | Farnesyl diphosphate synthase [EC:2.5.1.1 2.5.1.10] |
| Pentose phosphate pathway | | | | | | |
| Oxidative branch | | | | | | |
| rtg5491 | 2656 | 0.297504692 | <0.1 | K07404 | G | 6-phosphogluconolactonase [EC:3.1.1.31] |
| rtg994 | 695 | 0.586741311 | <0.01 | K01057 | G | 6-phosphogluconolactonase [EC:3.1.1.31] |
| rtg3203 | 190 | 0.93934475 | <0.01 | K00033 | G | 6-phosphogluconate dehydrogenase [EC:1.1.1.44 1.1.1.343] |
| Non-oxidative branch | | | | | | |
| rtg2879 | 3444 | 0.363202447 | ns | K00615 | G | Transketolase [EC:2.2.1.1] |
| rtg2881 | 307 | 0.806224211 | <0.01 | K00615 | G | Transketolase [EC:2.2.1.1] |
| rtg5013 | 13 | 1.94948339 | <0.01 | K01621 | G | Xylulose-5-phosphate/fructose-6-phosphate phosphoketolase [EC:4.1.2.9 4.1.2.22] |
| rtg5082 | 2674 | 0.27808639 | <0.1 | K01807 | G | Ribose 5-phosphate isomerase A [EC:5.3.1.6] |
| rtg11 | 5616 | 0.011283002 | ns | K01783 | G | Ribulose-phosphate 3-epimerase [EC:5.1.3.1] |
| rtg4731 | 1157 | 0.451830958 | <0.01 | K00616 | G | Transaldolase [EC:2.2.1.2] |
| rtg3728 | 564 | -0.769286911 | <0.01 | K15634;K22315 | G | Sedoheptulose-bisphosphatase [EC:3.1.3.37] |
| rtg1810 | 1269 | -0.554187819 | <0.1 | K00852 | G | Ribokinase [EC:2.7.1.15] |
| rtg1972 | 1920 | -0.458514409 | <0.05 | K01835 | G | Phosphoglucomutase [EC:5.4.2.2] |
| rtg2834 | 2125 | 0.426807962 | <0.01 | K01835 | G | Phosphoglucomutase [EC:5.4.2.2] |
| rtg3427 | 3269 | -0.381710813 | ns | K00948 | F | Ribose-phosphate pyrophosphokinase [EC:2.7.6.1] |
